# Molecular mechanism of translational stalling by inhibitory codon combinations and poly(A) tracts

**DOI:** 10.1101/755652

**Authors:** Petr Tesina, Laura N. Lessen, Robert Buschauer, Jingdong Cheng, Colin Chih-Chien Wu, Otto Berninghausen, Allen R. Buskirk, Thomas Becker, Roland Beckmann, Rachel Green

## Abstract

Inhibitory codon pairs and poly(A) tracts within the translated mRNA cause ribosome stalling and reduce protein output. The molecular mechanisms that drive these stalling events, however, are still unknown. Here, we use a combination of *in vitro* biochemistry, ribosome profiling, and cryo-EM to define molecular mechanisms that lead to these ribosome stalls. First, we use an *in vitro* reconstituted yeast translation system to demonstrate that inhibitory codon pairs slow elongation rates which are partially rescued by increased tRNA concentration or by an artificial tRNA not dependent on wobble base pairing. Ribosome profiling data extend these observations by revealing that paused ribosomes with empty A sites are enriched on these sequences. Cryo-EM structures of stalled ribosomes provide a structural explanation for the observed effects by showing decoding-incompatible conformations of mRNA in the A sites of all studied stall-inducing sequences. Interestingly, in the case of poly(A) tracts, the inhibitory conformation of the mRNA in the A site involves a nucleotide stacking array. Together, these data demonstrate novel mRNA-induced mechanisms of translational stalling in eukaryotic ribosomes.

## Introduction

Coding sequences for proteins in any genome (the open reading frames or ORFs) have evolved in the context of their full mRNA transcript to be expressed at the appropriate level. Interestingly, synonymous codon choice has been shown to have broad impacts on many aspects of translation including translational efficiency (Gingold & Pilpel, 2011, Tuller, Waldman et al., 2010), mRNA decay (Presnyak, Alhusaini et al., 2015a) and cotranslational protein folding (Pechmann & Frydman, 2013, Thanaraj & Argos, 1996). The effects on translational efficiency are primarily mediated through the competition of cognate and near-cognate tRNA interactions, as dictated by the pool of charged tRNAs available in the cell (Dana & Tuller, 2014, Elf, Nilsson et al., 2003, Gingold & Pilpel, 2011). Individual codons that are generally decoded by more abundant tRNAs and are associated with increased translation efficiency have been defined as “optimal” (Burgess-Brown, Sharma et al., 2008, dos Reis, Savva et al., 2004, Sharp & Li, 1987). Moreover, codon usage biases, codon context and interactions between adjacent codons have all been suggested to play a role in translational efficiency (Brule & Grayhack, 2017, Quax, Claassens et al., 2015), though their direct effects on elongation are still not fully understood.

A recent study in yeast defined a collection of 17 specific codon pairs that caused a substantial down-regulation in protein output (Gamble, Brule et al., 2016). For 12 of these pairs, the order of the codons within the pair was critical for the observed inhibition. Despite the diverse nature of these pairs, there were some shared features. First, the proline codon CCG and the arginine codon CGA appeared frequently in the collection of inhibitory pairs. The CCG codon is decoded by a G-U wobble base pair while the CGA codon is the sole codon in yeast decoded by an obligate I:A wobble pair (Letzring, Dean et al., 2010). Notably, while the previous study (Gamble et al., 2016) concluded that these inhibitory codon pairs likely impacted the decoding step of elongation, there was little understanding of the molecular basis for these events.

Besides inhibitory codon pairs, poly(A) tracts represent perhaps the most abundant and potent stall-inducing mRNA sequence in eukarya (reviewed in (Arthur & Djuranovic, 2018). Translation of poly(A) sequences commonly occurs when ribosomes encounter an abnormal (premature) polyadenylation event within the ORF or when ribosomes read through a stop codon. Premature polyadenylation alone occurs in approximately 1% of yeast and human transcripts, highlighting the importance of this mechanism (Frischmeyer, van Hoof et al., 2002, Ozsolak, Kapranov et al., 2010). While translation of poly(A) tracts initially results in the synthesis of poly-lysine, long poly(A) tracts subsequently trigger quality control pathways that contribute to overall protein homeostasis (Brandman & Hegde, 2016, Joazeiro, 2019). The earliest studies suggested that this stalling was caused by electrostatic interactions between the poly-basic nascent chain and the peptide exit tunnel of the ribosome (Lu & Deutsch, 2008). However, there are several lines of evidence suggesting that the stalling mechanism of poly(A) tracts is more complex. Interestingly, as few as two consecutive AAA codons were shown to cause ribosome sliding during translation in *E. coli* (Koutmou, Schuller et al., 2015). Moreover, the identity of the basic residue-encoding codon is of particular importance for efficient stalling, as the CGA arginine-encoding codon is most potent in yeast (Letzring, Wolf et al., 2013), and AAA codons are more potent than AAG lysine-encoding codons at inducing translational stalling (Arthur, Pavlovic-Djuranovic et al., 2015, Koutmou et al., 2015).

### Inhibitory codon pairs slow elongation *in vitro*

To examine the impact of inhibitory codon pairs on translation elongation *in vitro*, we selected pairs that most potently reduced GFP expression in the *in vivo* experiments and those that contained codons which appeared in multiple inhibitory pairs (Fig 1A) (Gamble et al., 2016). The strongest candidates were CGA-CGA and CGA-CCG encoding Arg-Arg and Arg-Pro, respectively. The arginine codon CGA is decoded by ICG tRNA^Arg^ where inosine forms a unique purine-purine I:A wobble pair. The proline codon CCG is found in many inhibitory codon pairs, likely because it is decoded by tRNA using a G-U wobble pair, UGG tRNA^Pro^ (Fig 1A). The prevalence of and dependency on wobble base-pairing in inhibitory codon pairs led Grayhack and co-workers to conclude that elongation is blocked by non-optimal codon-anticodon pairing at neighboring sites on the ribosome (i.e. the P and A sites). Furthermore, they showed that for these codon pairs, the order of the codons in the pair is critical; the reverse pair has little to no effect on protein output.

**Figure 1.**
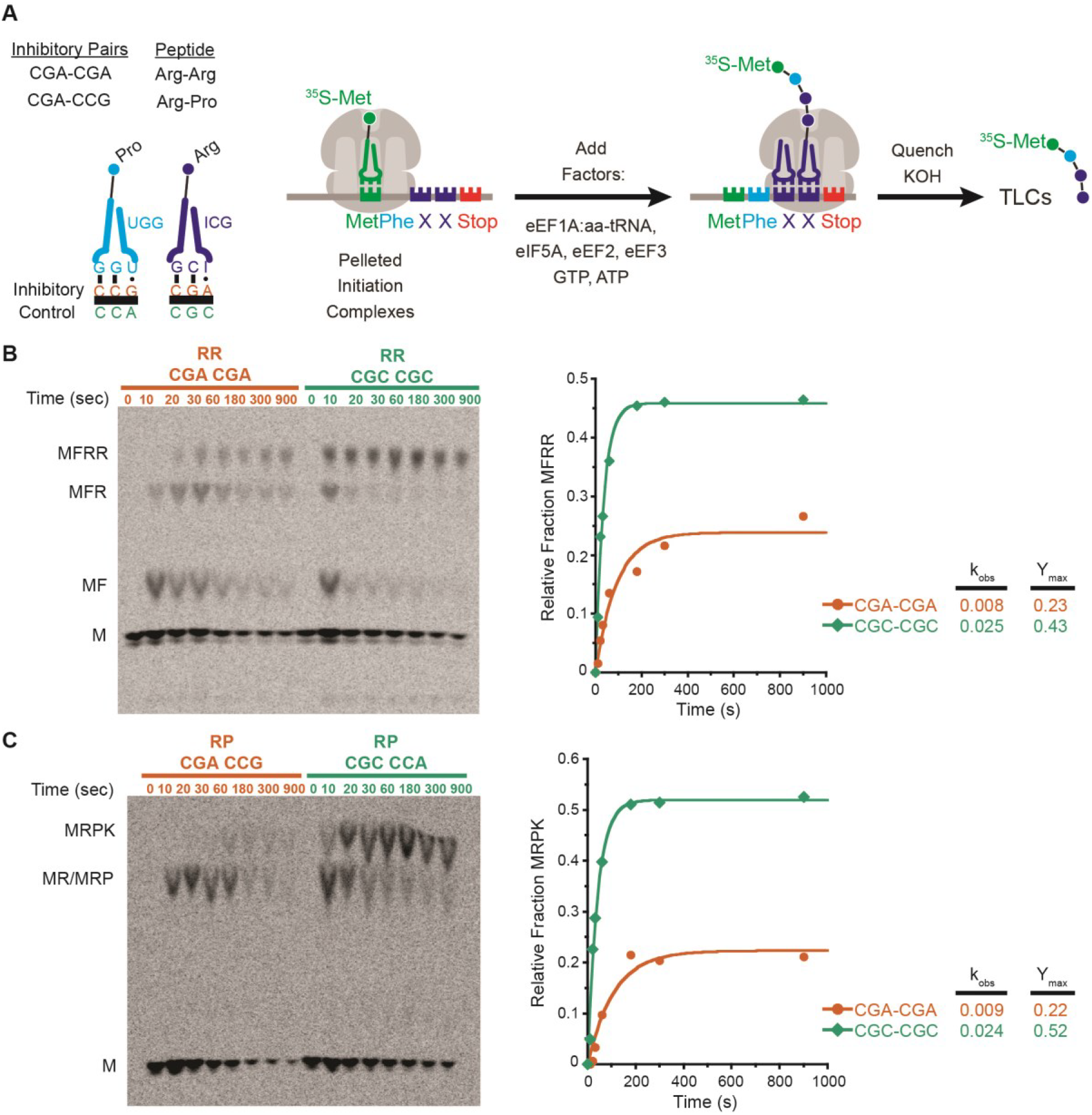
Inhibitory codon pairs slow elongation *in vitro*. (**A**) Inhibitory pairs showing the inhibitory mRNA codons (red) and the optimal codons (green). Schematic representation of the *in vitro* elongation reactions performed using the reconstituted yeast translation system. (**B**) Representative eTLCs (left) and corresponding elongation kinetics (right) for the CGA-CGA inhibitory pair (red) and the CGC optimal pair (green). (**C**) Representative eTLCs (left) and corresponding elongation kinetics (right) for the CGA-CCG inhibitory pair (red) and the CGC-CCA optimal pair (green). Product formation in (**C**) and (**D**) are normalized to the fraction of Met ICs that form Met-Puro when reacted with puromycin (Fig EV1A).

All of the inhibitory sequences described above result in partial or complete translational stalling *in vivo*. Considerable attention has been paid to the molecular consequences of the translating ribosomes encountering such mRNA sequences (i.e. the downstream quality control events that are triggered). In particular, recent work has suggested that ribosomal collisions with the leading, stalled ribosome are a key event that triggers the quality control responses that include decay of the mRNA (“No Go Decay” or NGD) and the nascent peptide (Ribosome-associated Quality Control or RQC) (Ikeuchi, Tesina et al., 2019b, Juszkiewicz, Chandrasekaran et al., 2018b, Simms, Yan et al., 2017). However, there has been little characterization of the molecular events on the ribosome that lead to such dramatic outcomes.

Here we use a yeast *in vitro* reconstituted biochemical system to directly measure the rates of translation elongation that might be impacted by inhibitory codon pairs and poly(A) tracts. Use of this *in vitro* system allows for ready manipulation of mRNA coding sequence, tRNA identity and concentration, as well as ribosome composition to reveal defects in the individual steps of translation elongation. Together with high-resolution ribosome profiling, our results reveal clear defects in the decoding step as the primary determinant of ribosomal stalling on these inhibitory mRNA sequences. Cryo-EM structures of ribosome complexes stalled at these mRNA sequences reveal detailed insight into the molecular basis for the translational stalling. Importantly, we observe decoding-incompetent conformations of mRNA in the A sites of all stall-inducing sequences that we studied, thus readily explaining the biochemically-defined decoding defects. Moreover, structural characterization of poly(A) stalled disomes reveals a novel disome conformation with both ribosomes in the POST translocation state, suggesting a role for ribosome collisions in promoting frameshifting. Taken together, our data reveal an mRNA-induced translational stalling mechanism of eukaryotic ribosomes.

To monitor synthesis of tetrapeptides containing these inhibitory codon pairs, we employed an *in vitro* reconstituted yeast translation system (Eyler & Green, 2011, Schuller, Wu et al., 2017). Initiation complexes (ICs) were assembled using ribosome subunits, [^35^S]-Met-tRNA^iMet^, and mRNAs containing an AUG codon, the codon pair of interest, and an additional codon encoding Phe or Lys before or after the pair to enhance visualization of the products by electrophoretic thin-layer chromatography (eTLC). Following purification, each IC was treated with puromycin (Pm) to release the nascent chain and determine the fraction of bound [^35^S]-Met-tRNA^iMet^ that forms Met-Pm. Puromycin reacts with peptidyl-tRNA bound to the ribosome when the peptidyl-transferase center (PTC) of the large subunit is accessible and releases the polypeptide chain as peptidyl-puromycin. As such, this assay reports on the overall competence and conformation of the peptidyl-transferase center of the ICs. We consistently observed that ICs formed with the different mRNA transcripts formed Met-Pm products to a similar extent (Fig EV1A). Therefore, differences in the amount of peptide produced using ICs containing different mRNA templates were not due to the efficiency of IC formation or to the differential ability of the programmed ribosome to make peptide bonds.

For elongation reactions, the desired tRNAs were purified from bulk tRNA using biotinylated oligonucleotides (Yokogawa, Kitamura et al., 2010), charged with the corresponding aminoacyl-tRNA synthetase, and the aminoacyl-tRNAs were pre-incubated with eEF1A and GTP to form ternary complexes. Ternary complexes were then mixed with purified ICs and elongation factors eEF2, eEF3, and eIF5A. Peptide formation was monitored by quenching time points of the reactions in KOH and resolving the formed products by eTLC (Fig 1A). The initial experiments were performed with ribosome complexes at ~2 nM and aa-tRNAs at ~12 nM, where both binding and catalysis contribute to the observed rate (i.e. k_cat_/K_m_ conditions). For each inhibitory codon pair a control “optimal” IC was prepared where the non-optimal codons were replaced by synonymous codons that are decoded by the same tRNA, but without wobble base pairing. For example, the optimal codon CGC was used as a control for CGA because it is decoded by the same ICG tRNA^Arg^ via a pyrimidine-purine C:I pair with a standard Watson-Crick geometry (Murphy & Ramakrishnan, 2004) instead of a purine-purine (A:I) wobble base pair (Fig 1A).

Visual examination of the reaction profiles for the inhibitory CGA-CGA codon pair (in red) relative to the optimal CGC-CGC codon pair (in green) reveals a clear defect in elongation (Fig 1B). First, the inhibitory Arg-Arg pair exhibits a significantly lower endpoint, with ~25% of the radiolabeled Met forming the final tetrapeptide product, MFRR, compared with ~45% for the optimal Arg-Arg sequence. In a similar fashion, there are clear elongation defects for the inhibitory Arg-Pro, CGA-CCG codon pair (in red) compared to the optimal CGC-CCA codon pair (in green) (Fig 1C); the inhibitory Arg-Pro pair has only ~20% of the radiolabeled Met forming the final tetrapeptide product, MRPK, compared to ~50% for the optimal Arg-Pro sequence. As endpoint defects often suggest the existence of an off pathway reaction, we asked whether there were high levels of peptidyl-tRNA drop-off during elongation for the Arg-Arg or Arg-Pro reactions that might explain the observed defects. However, when we directly tested this possibility using an assay involving peptidyl hydrolase (Pth) that acts only on tRNAs not bound to the ribosome, we saw no evidence for drop off with any of the complexes (Fig EV1B, EV1C) (Schuller et al., 2017, Shoemaker, Eyler et al., 2010).

In addition to the endpoint defects, we also observe a reduced rate of formation of the final peptide product for the complexes encoding both the Arg-Arg and Arg-Pro pairs; in each case, the observed rates were about three-fold slower than those of their optimal counterparts (Fig 1B, C). For the Arg-Arg pair, where MFR and MFRR can be separately resolved, we see a substantial build-up of MFR intermediate peptide relative to the CGC-CGC dicodon control (Figs 1B and EV1D). Quantification of both products (MFR and MFRR) of this inhibitory pair as well as elongation on a single arginine message (MFR) indicate that elongation through the first CGA codon is slightly slow, but that the subsequent elongation through the second CGA codon is the major inhibitory step (Figs EV1E, EV1F).

Together, these data reveal *in vitro* defects in elongation reactions on the ribosome resulting from two distinct inhibitory codon pairs. These observations provide strong evidence that the initially observed effects *in vivo* (Gamble et al., 2016) reflect defects intrinsic to ribosome function rather than resulting from mRNA decay or other downstream cellular events.

### Multiple defects in decoding caused by codon pairs

Assuming that one likely cause of elongation slow-down may be defects in decoding, we asked if the inhibition arises from simple defects in the energetics of tRNA binding (a second order event) or instead from more downstream defects (i.e. in first order events) that follow including GTPase activation and accommodation (Gromadski & Rodnina, 2004, Zaher & Green, 2009). As the initial *in vitro* experiments were performed in a k_cat_/K_m_ regime, we repeated the elongation assays at 10-fold higher ternary complex concentrations. For Arg-Arg, we see an approximately 2-fold rescue of the rate of the reaction with higher tRNA concentrations for the inhibitory pair (CGA-CGA) with only very modest changes in the rate of the reaction for the optimal pair (CGC-CGC) (Fig 2A, left). Similarly, for the Arg-Pro combination, we see an approximately 4-fold increase in the rate of the reaction with higher tRNA concentrations for the inhibitory pair (CGA-CCG) with only a modest, maximally 1.5-fold increase, for the optimal pair (CGC-CCA) (Fig 2A, right). These results suggest that tRNA binding contributes in part to the observed defects seen for the inhibitory pairs. Importantly, however, we observe that for both codon pairs (CGA-CGA and CGA-CCG), the endpoint defects are not overcome at high tRNA concentrations (Fig 2B). These latter data strongly suggest that a certain fraction of the complexes is unable to elongate independent of saturating levels of aminoacyl-tRNA substrate.

**Figure 2.**
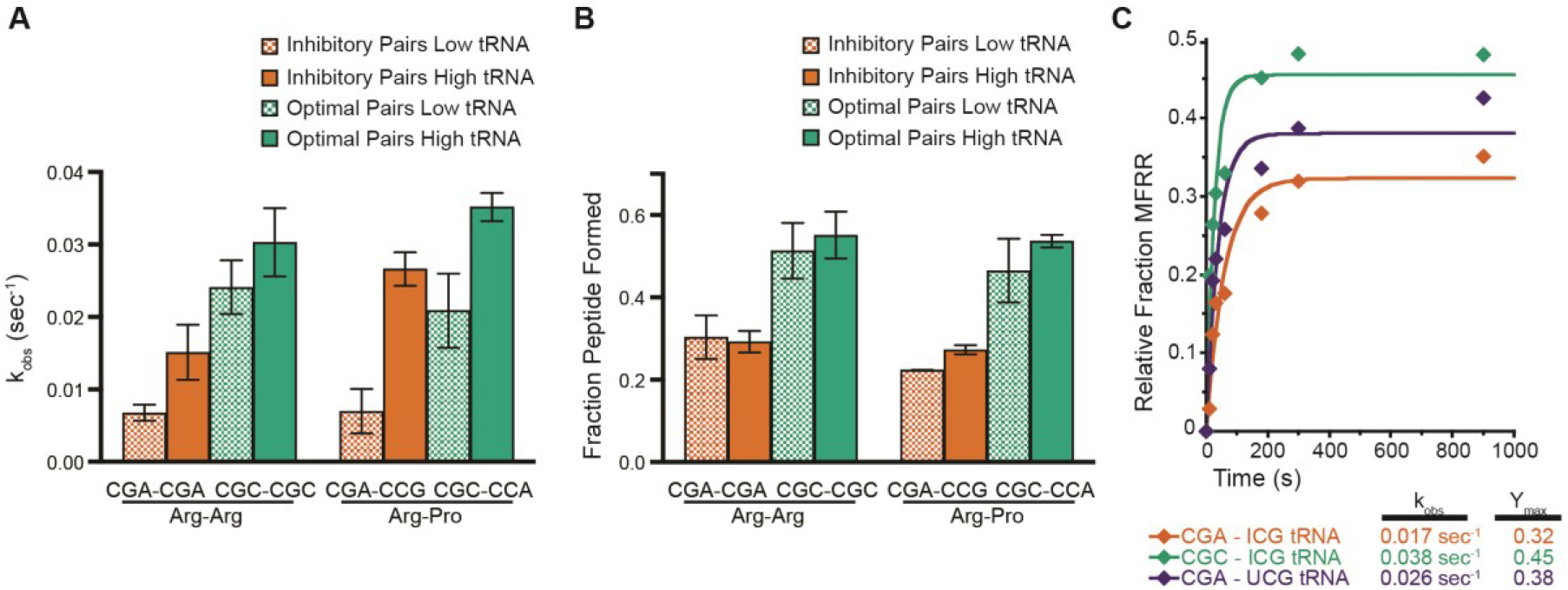
Effects of tRNA concentration on elongation rates and endpoints. **(A)** Comparison of observed rates of elongation for inhibitory pairs (red) and their optimal controls (green) at limiting tRNA concentrations (hatched bars) and saturating tRNA concentrations (solid bars). **(B)** Comparison of total peptide formation for inhibitory pairs (red) and their optimal controls (green) at limiting tRNA concentrations (hatched bars) and saturating tRNA concentrations (solid bars). **(C)** Elongation kinetics for the CGA-CGA inhibitory codon pair with the native ICG tRNA^Arg^ (red) or the non-native UCG tRNA^Arg^ (purple) and for the CGC-CGC optimal control pair with the native ICG tRNA^Arg^ (green). Error bars represent standard deviations calculated from at least three experimental replicates.

Given the unusual nature of the I:A wobble base pair found in the P site after incorporation of the first Arg in the codon pair, we also wondered whether the substantial defects that we observed might be rescued with the use of a non-natural, exact match UCG tRNA^Arg^ as shown *in vivo* in the previous study (Gamble et al., 2016). We expressed the non-natural tRNA^Arg^ on a CEN plasmid in yeast and purified it as above using a biotinylated oligonucleotide. In elongation reactions performed under k_cat_ conditions (high tRNA concentrations), this non-natural tRNA did partially rescue the endpoint defects in the elongation reaction associated with the CGA-CGA codon pair (Fig 2C); these data suggest that the unusual I:A pairing in the P site at least partially contributes to the endpoint defects associated with these inhibitory codon pairs.

### Increased 21 nt RPFs on inhibitory pairs indicate an empty ribosomal A site

To further investigate the molecular mechanisms of inhibition underlying the inhibitory codon pairs, we turned to high-resolution ribosome profiling (Wu, Zinshteyn et al., 2019). We recently reported that ribosome profiling using a cocktail of elongation inhibitors can trap ribosomes in their different functional states, distinguished by the size of ribosome protected footprints (RPFs). For example, when cycloheximide (CHX) and tigecycline (TIG) are added to yeast lysates to prevent ribosomes from translating post cell lysis, RPFs that are 21 nucleotides (nts) in length correspond to ribosomes in a “classical” or POST state waiting to decode the next aminoacyl-tRNA while RPFs that are 28 nts in length correspond primarily to ribosomes trapped in a “rotated” or PRE state (Wu et al., 2019). Building on an earlier study that showed an enrichment in ribosome density when the 17 inhibitory codon pairs are aligned (Gamble et al., 2016, Matsuo, Ikeuchi et al., 2017), we generated libraries using CHX and TIG to better distinguish the functional state of the paused ribosomes. In the plot shown in Fig 3A, the average ribosome density on 17 inhibitory codon pairs (with the first codon in the P site and second in the A site) is shown as a function of the RPF length on the Y-axis. We observe that while the density of 28 nt RFPs is fairly constant across this region, there is a large accumulation of 21 nt RFPs at the A site codon (Fig 3A). These data indicate that for these 17 inhibitory pairs, elongation inhibition is likely caused by slow decoding of the second codon of the inhibitory pair, resulting in an empty A site that yields shorter footprints.

**Figure 3.**
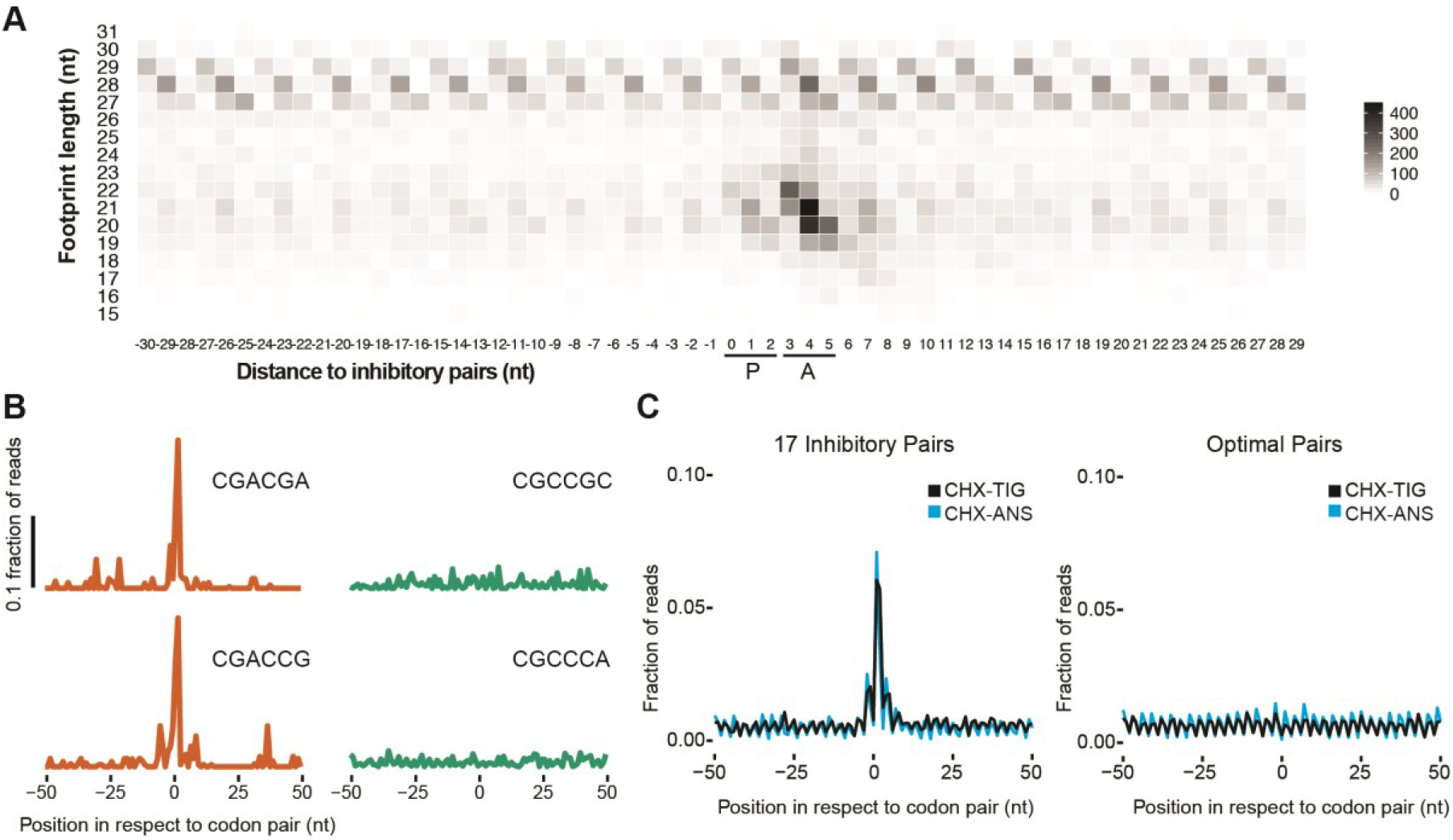
Increased 21 nt RPFs on inhibitory pairs indicate an empty ribosomal A site. (**A**) Meta-analysis of footprint size of all 17 inhibitory pairs identified by Grayhack and coworkers (Gamble et al., 2016), aligning the first codon of the pair in the ribosomal P site. (**B**) Metacodon analysis of 21 nt RPFs centered at the first codon of each inhibitory pair (red) compared to their corresponding optimal pair (green). (**C**) Comparison of 21 nt RPFs aligned at all 17 inhibitory codons from libraries made with CHX/ANS (blue) and CHX/TIG (black) (left) to their corresponding optimal pairs with the same antibiotic combination (right).

We can also look individually at the representative codon pairs studied above (CGA-CGA and CGA-CCG) and we see significant accumulation of 21 nt RPFs in the A site relative to the amount observed for their optimal counterparts (red vs. green) (Fig 3B). These data provide direct evidence that elongation inhibition on these codon pairs results from slow decoding of the second codon of the inhibitory pair.

We also considered the possibility that for the inhibitory codon pairs, tRNAs are accommodated but fail to undergo peptidyl transfer, perhaps because of a misalignment in the active site of the 60S subunit. We observed previously that the addition of anisomycin (ANS), a peptidyl-transferase inhibitor, together with CHX, blocks bound tRNAs from forming peptide bonds such that they eventually fall out of the A site; in these libraries, 21 nt RPFs represent two different ribosome populations, those in a pre-accommodation and a pre-peptidyl transfer state. Indeed, in samples prepared with CHX/ANS, we observe more 21 nt RPFs at peptide motifs known to undergo slow peptidyl transfer (Schuller et al., 2017) relative to those motifs or codons enriched in the CHX/TIG samples (Fig EV2A). If the chemistry of peptide-bond formation were slow for the inhibitory codon pairs, we would expect to see an increase of 21 nt RPFs at these sites in the CHX/ANS library relative to the CHX/TIG library. Instead, we see the same level of enrichment of 21 nt RPFs at these sites (Fig 3C, left), arguing that the limiting step for the inhibitory base pairs is not peptide bond formation. These findings are consistent with the hypothesis that certain wobble pairs impact the decoding center in the 40S subunit, affecting decoding or accommodation, rather than activities in the peptidyl-transferase center of the large subunit. For the optimal codon pairs for these same amino acid sequences, no pauses are seen in either sample indicating that the pausing at inhibitory codons is due to the codon/tRNA pairing in the A site rather than to the amino acid sequence (Fig 3C, right).

### Loss of the ribosomal protein Asc1 inhibits elongation

Several studies in yeast using iterated CGA codons to induce ribosome stalling have shown that the loss of the ribosomal protein Asc1 enables ribosomes to read through these inhibitory sequences (Letzring et al., 2013, Wang, Zhou et al., 2018, Wolf & Grayhack, 2015); these data suggest that Asc1 is somehow involved either in facilitating proper decoding or in sensing and stabilizing stalled ribosomes. We asked what role Asc1 plays in the elongation of CGA codon pairs using our *in vitro* system. We first prepared ribosomes from an Asc1 deletion strain and produced initiation complexes programmed with either non-optimal (CGA-CGA) or optimal (CGC-CGC) MFRR mRNAs as before and compared their elongation reactions. Initiation complex formation and the puromycin reactivity of these complexes was indistinguishable from that of complexes formed with wild-type ribosomes (Fig EV3A). Elongation reactions were then performed as described above using ICG tRNA^Arg^ to decode the Arg codons in both mRNAs. We see that for both the inhibitory and optimal di-codon pair complexes, ribosomes lacking Asc1 elongate more slowly and reach a lower elongation endpoint (Fig 4A). These data suggest that ribosomes lacking Asc1 have general defects in elongation. Elongation reactions with ICs lacking Asc1 for the CGA-CCG, Arg-Pro pair show similar defects (Fig EV3B). We also performed high-resolution ribosome profiling in an *asc1Δ* strain using CHX/TIG for the preparation as above (Wu et al., 2019). In this analysis, we observe a genome-wide increase of 21 nt RPFs, consistent with the idea that ribosomes lacking Asc1 broadly struggle with the tRNA decoding step of translation elongation (Fig 4B). Moreover, when we specifically look at the pausing signature of ribosomes at the 17 inhibitory codon pairs, we see that the CGA-CCG and CGA-CGA codon pairs show the largest enrichment in 21 nt RPFs in the *asc1* deletion strain compared to the wild-type strain (Fig 4C). Together, these data provide support for the idea that the ribosomal protein Asc1 makes important contributions to the tRNA selection step of translation elongation

**Figure 4.**
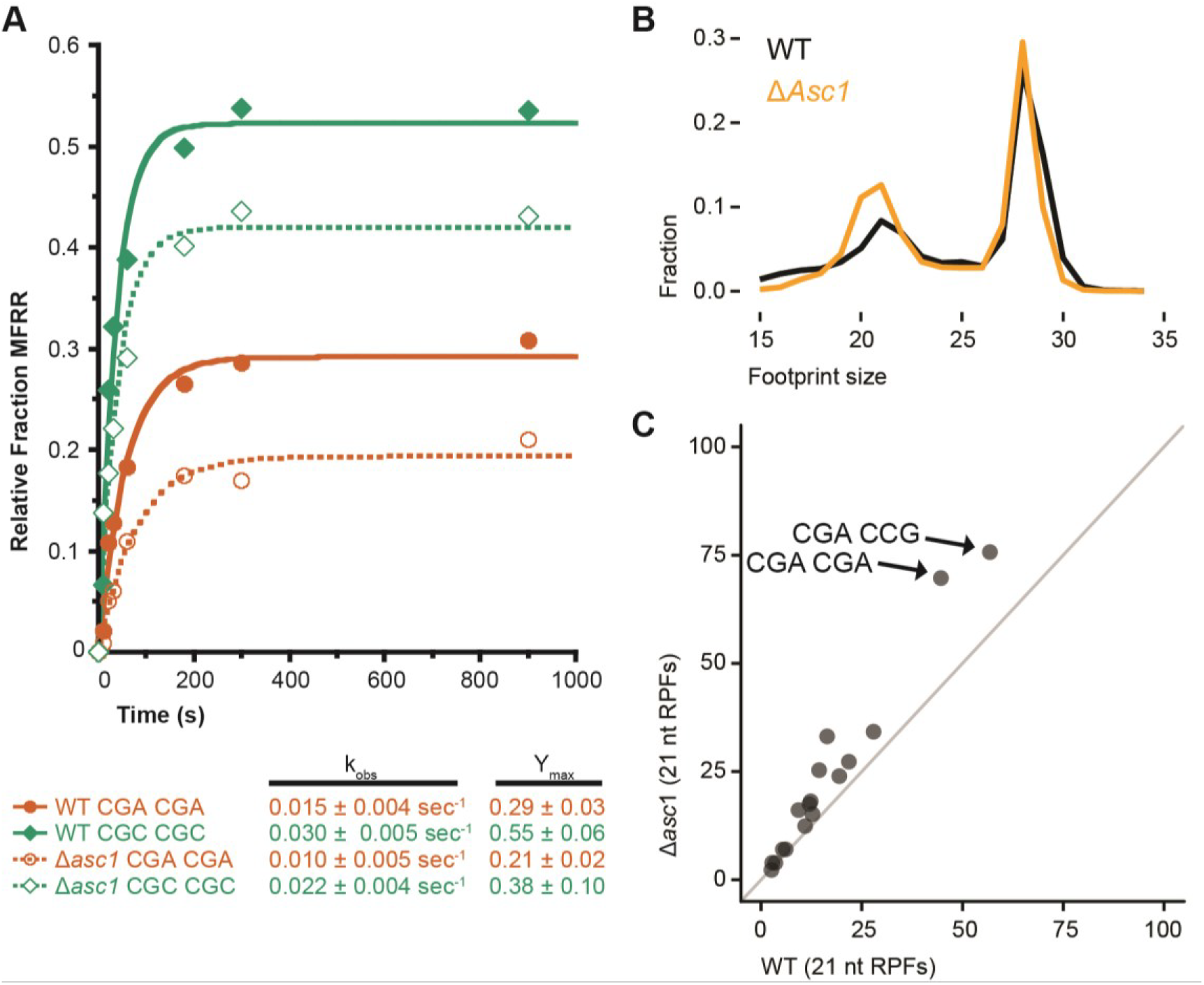
Loss of the ribosomal protein Asc1 inhibits elongation. (**A**) Elongation kinetics for the CGA-CGA inhibitory codon pair (red) and the CGC-CGC optimal control (green) from WT ribosomes (solid) or the *asc1Δ* strain ribosomes (dashed) at saturating tRNA concentrations. Average observed rates and elongation endpoints from three or more replicate experiments shown below the graph. (**B**) Size distributions of ribosome footprints for WT cells (black) and *asc1Δ* cells (gold). (**C**) Scatter plot of ribosome occupancies for 21 nt RPFs at the 17 inhibitory codon pairs (Gamble et al., 2016) comparing WT cells to *asc1Δ* cells with the two inhibitory codon pairs further investigated in this study labeled.

### Decoding-incompatible mRNA conformation causes to inhibitory codon pair-mediated stalling

To investigate the molecular basis of the inhibitory codon pairs involving the problematic CGA codon, we turned to structural studies of complexes stalled at CGA-CCG and CGA-CGA codon pairs. We used a yeast cell-free *in vitro* translation system in which we translated mRNA reporters containing the CGA-CCG or CGA-CGA inhibitory codon pairs. Translation extracts were prepared from yeast cells lacking Ski2p, a component of the 3’-5’ mRNA decay system, to enhance mRNA stability. Both mRNA reporters contained sequences coding for an N-terminally His_8_-HA-tagged truncated uL4 (Knorr, Schmidt et al., 2019) followed by the stalling (CGA-CCG)_2_ or (CGA-CGA)_2_ codon pairs (Appendix Figs S1A and S2A). To avoid capturing read-through products, the stalling sequences were followed by three UAA(A) stop codon quadruplets, one in each reading frame, which would lead to termination upon read-through. Ribosome nascent chain complexes (RNCs) were affinity purified using magnetic beads, separated on a sucrose density gradient and the 80S fractions were subjected to cryo-EM (Appendix Figs S1 and S2).

Classification of ribosomal particles for both stalling sequences (CGA-CCG and CGA-CGA) revealed the most abundant classes to be programmed ribosomes in the post-translocation state (POST state) with tRNAs in the P/P and E/E state, but not in the A site (Appendix Figs S3 and S4). The structure of the CGA-CCG stalled ribosome was reconstructed to an average resolution of 2.6 Å while the CGA-CGA stalled ribosome was reconstructed to an average resolution of 3.2 Å (Fig EV4B and C). To compare these structures on a molecular level with a canonical A site tRNA decoding situation, we refined our previously produced structure of cycloheximide-stalled ribosomes in the pre-translocation state (PRE state) with A/A and P/P tRNAs to 3.1 Å with focus on the mRNA decoding in the A site (Figs 5A and EV4A) (Buschauer et al. 2019). Molecular models were built and refined for all structures allowing for an in-depth analysis (Fig 5A-I). Structural analysis of the CGA-CCG and the CGA-CGA stalled RNCs revealed no perturbations of the peptidyl-transferase center (PTC), in agreement with the puromycin reactivity of these stalled ribosomes (Appendix Fig S5). On the other hand, we saw a strikingly unusual conformation of the mRNA in the A site of these structures when compared with the canonical decoding situation (Fig 5A-I).

**Figure 5:**
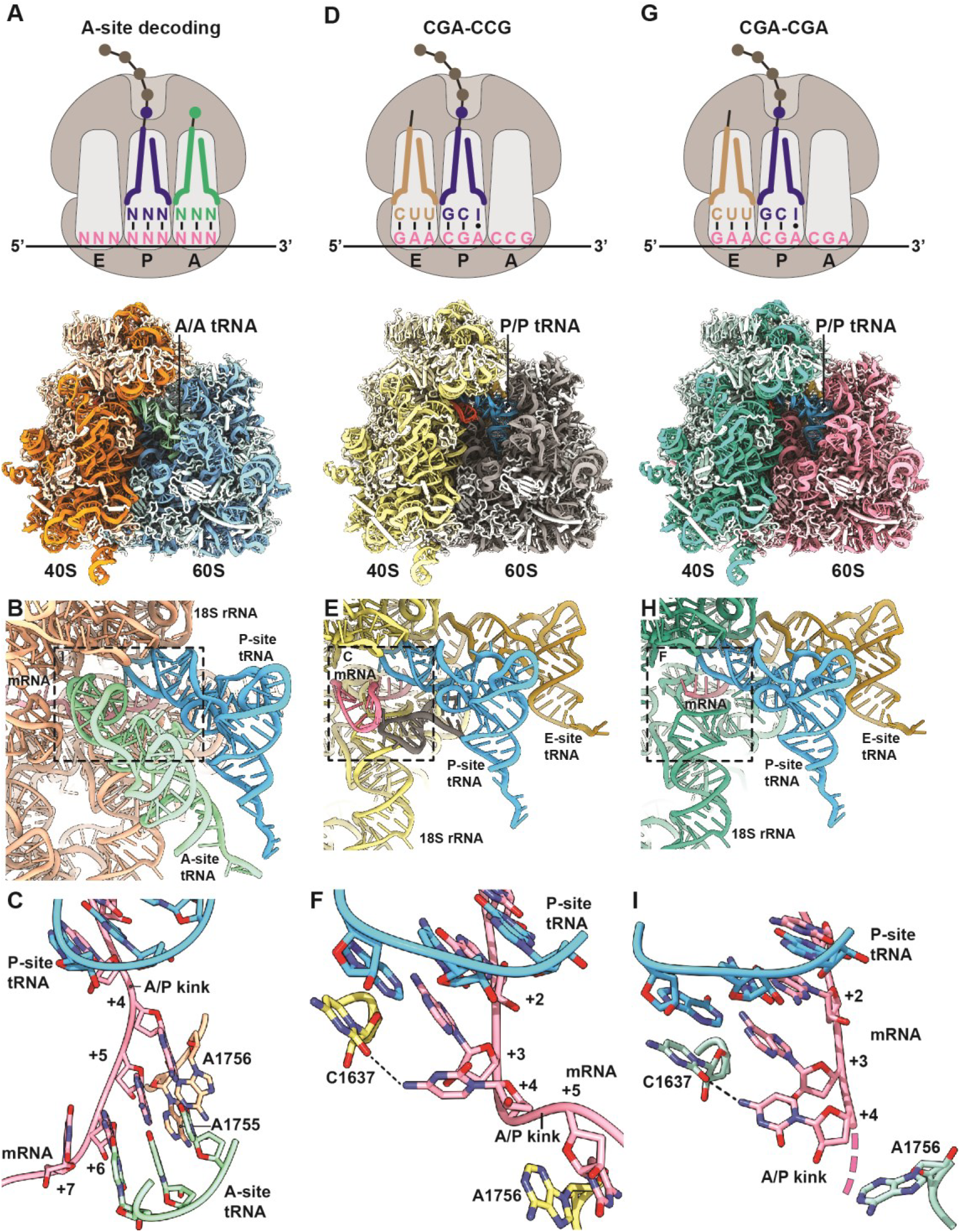
CGA-CCG and CGA-CGA induce stalling through decoding-incompatible mRNA conformations in the A site. (**A-C**) Cryo-EM structural characterization of the pre-state RNC with A site tRNA in the decoding center. (**A**) Schematic representation of the decoding situation (top) and molecular model for the pre-state RNC with A site tRNA in the decoding center. (**B**) General overview of the A, P and E sites with A/A and P/P tRNAs and mRNA. (**C**) Detailed view of the mRNA in the A site using sticks model with cartoon phosphate backbone representation. The 18S rRNA bases A1755 and A1756 recognize the minor groove of A site tRNA – mRNA interaction during tRNA decoding. (**D-F**) Cryo-EM structural characterization of the CGA-CCG stalled RNC. (**D**) Schematic representation of the stalling situation (top) and molecular model of the CGA-CCG stalled RNC (bottom). (**E**) General overview of the A, P, and E sites with P/P and E/E tRNAs and mRNA. (**F**) Detailed view of the mRNA in the A site using sticks model with cartoon phosphate backbone representation. The mRNA positions +2 to +5 and their interactions are shown. The C +4 is flipped by approximately 95° degrees towards the wobble A:I base pair in the P site and stabilized by interaction with the C1637 of the 18S rRNA helix 44. The C +5 is stabilized by stacking interaction with the A1756 of the 18S rRNA which normally recognizes the minor groove of A site tRNA – mRNA interaction during decoding (**L**). **(G-I)** Cryo-EM structural characterization of the CGA-CGA stalled RNC. (**G**) Schematic representation of the stalling situation (top) and molecular model of the CGA-CGA stalled RNC (bottom). (**H**) General overview of the A, P and E sites with P/P and E/E tRNAs and mRNA. (**I**) Detailed view of the mRNA in the A site as in (E). Downstream mRNA is indicated by the dotted line. Note the rotation of the C+4 base compared to the CGA-CCG mRNA.

The most striking mRNA structure is formed on the CGA-CCG reporter mRNA. In our 2.6 Å map, we can clearly identify the CGA-codon in the P site and the anticodon of ICG tRNA^Arg^ making standard Watson-Crick interactions as observed before (Schmidt, Kowalinski et al., 2016) at the first two positions of the codon and a purine:purine A:I base pair at the wobble position (Fig 5E). However, the first nucleotide in the A site (the C+4 of the CCG codon) is found in an unusual conformation that is well defined by the cryo-EM density (Fig EV5A). Compared to the control canonical decoding situation (Fig 5A-C), C+4 is flipped by approximately 95° degrees towards the wobble A:I base pair in the P site. Stabilization of C+4 in this position appears to be facilitated by an H-bond formed with C1637 of 18S rRNA helix 44 (C1400 in *E. coli*) which stacks on the I of the ICG tRNA^Arg^ in the P site (Fig 5F). Compared to the canonical decoding situation, accommodation of the purine:purine A:I wobble base pair at position +3 shifts the mRNA backbone by 2.6 Å at the phosphate linking +3 and +4, thus forcing the general path of the downstream mRNA into an unusual direction (Fig EV5B and C). Importantly, this alteration in the mRNA structure moves the crucial A/P kink to occur between positions +4 and +5 (Fig 5F). The A/P kink, normally positioned between positions +3 and +4, was shown to be crucial for A site interaction and proofreading activity, especially for difficult-to-decode near cognate tRNAs (Keedy, Thomas et al., 2018). In the flipped-out position seen here, the C+4 seems unlikely to be engaged by a canonical codon:anticodon interaction with the incoming aminoacyl-tRNA (Fig 5C). This rearrangement of the mRNA itself could explain the previously proposed communication between ribosomal P and A sites (Gamble et al., 2016).

Moreover, following C+4, the mRNA folds into a stable mRNA hairpin structure that directly occludes tRNA binding in the A site. In the hairpin, the C+5 base pairs with G+12 and the G+6 base pairs with C+11, while nucleotides C+7 – C+10 form a rather flexible tetraloop at the tip of the hairpin (Fig EV5D and E). Interestingly, this structure is stabilized by A1756 (A1493 in *E. coli)* of the 18S rRNA which flips out of helix h44 as well as the rearranged A2256 (A1913 in *E. coli*) of the 25S rRNA helix 69. Normally, A2256 forms a dynamic inter-subunit bridge 2A by intercalating into the 18S rRNA helix 44. However, to support the observed mRNA secondary structure formation, A2256 rotates by 101 degrees and stacks with C+7 of the mRNA (Fig EV5E). Taken together, this structure rationalizes how accommodation of the UGG-tRNA^Pro^ in the A site on the CGA-CCG inhibitory dicodon is prevented: i) by positioning of C+4 in a conformation incompatible with decoding, ii) by shifting the crucial mRNA A/P kink one position downstream and iii) by sterically blocking the tRNA binding site with an mRNA secondary structure.

Analogous to the CGA-CCG situation, we saw a specific inhibitory conformation of C+4 in the CGA-CGA mRNA cryo-EM structure (Fig 5G-I). Again, well supported by cryo-EM density, the conformation of C+4 is essentially the same as observed for the CGA-CCG reporter, with an 84° rotation of the cytosine base (Figs 5I and EV5F). After position +4, however, the mRNA density is weak and does not allow for reliable model building. These observations suggest a more flexible conformation of downstream mRNA in this structure. Nonetheless, the general path of mRNA seems to be shifted in the same direction as seen for the CGA-CCG case and the A/P kink in mRNA is also dislocated downstream as it cannot be observed between positions +3 and +4 (Figs 5I and EV5C). Taken together, these two structures show how rearrangement of the mRNA induced by the wobble-decoded CGA codon in the P site causes perturbations in the A site that disfavor decoding.

### Decoding-incompatible mRNA conformation contributes to poly(A) tract-mediated stalling

Next, we wondered whether the CGA-dependent codon pair stalling mechanism is structurally related to poly(A)-mediated stalling. First, using our *in vitro* system, we see slower elongation on a MFK_5_ AAA IC as compared to an AAG IC, consistent with earlier observations in *E. coli* (Koutmou et al., 2015). Despite resolution limitations of eTLC with multiple lysines, when we compare the earliest time points for AAA complexes with those for AAG complexes, the AAA complexes have only elongated to MF and MFK, whereas the AAG complexes are already making MK_2_ and larger products as indicated by the fast running smear (Fig EV6). These data are consistent with earlier reports documenting differences in elongation on iterated AAA relative to AAG lysine codons in other systems (Arthur et al., 2015, Koutmou et al., 2015).

For cryo-EM, we used an analogous approach to that used for CGA-dependent codon pair-mediated stalling with a modified mRNA reporter comprising a 49 nucleotide long poly(A) tract (Appendix Fig S6). As for both inhibitory codon pairs (CGA-CCG and CGA-CGA) discussed above, classification of poly(A) stalled ribosomal particles revealed that a majority (78%) of programmed particles are in the POST state without A-site tRNA (Appendix Fig S7). We reconstructed the poly(A)-stalled ribosome structure to an overall resolution of 3.1 Å, which allowed for building and refinement of a molecular model (Fig 6A and B).

**Figure 6:**
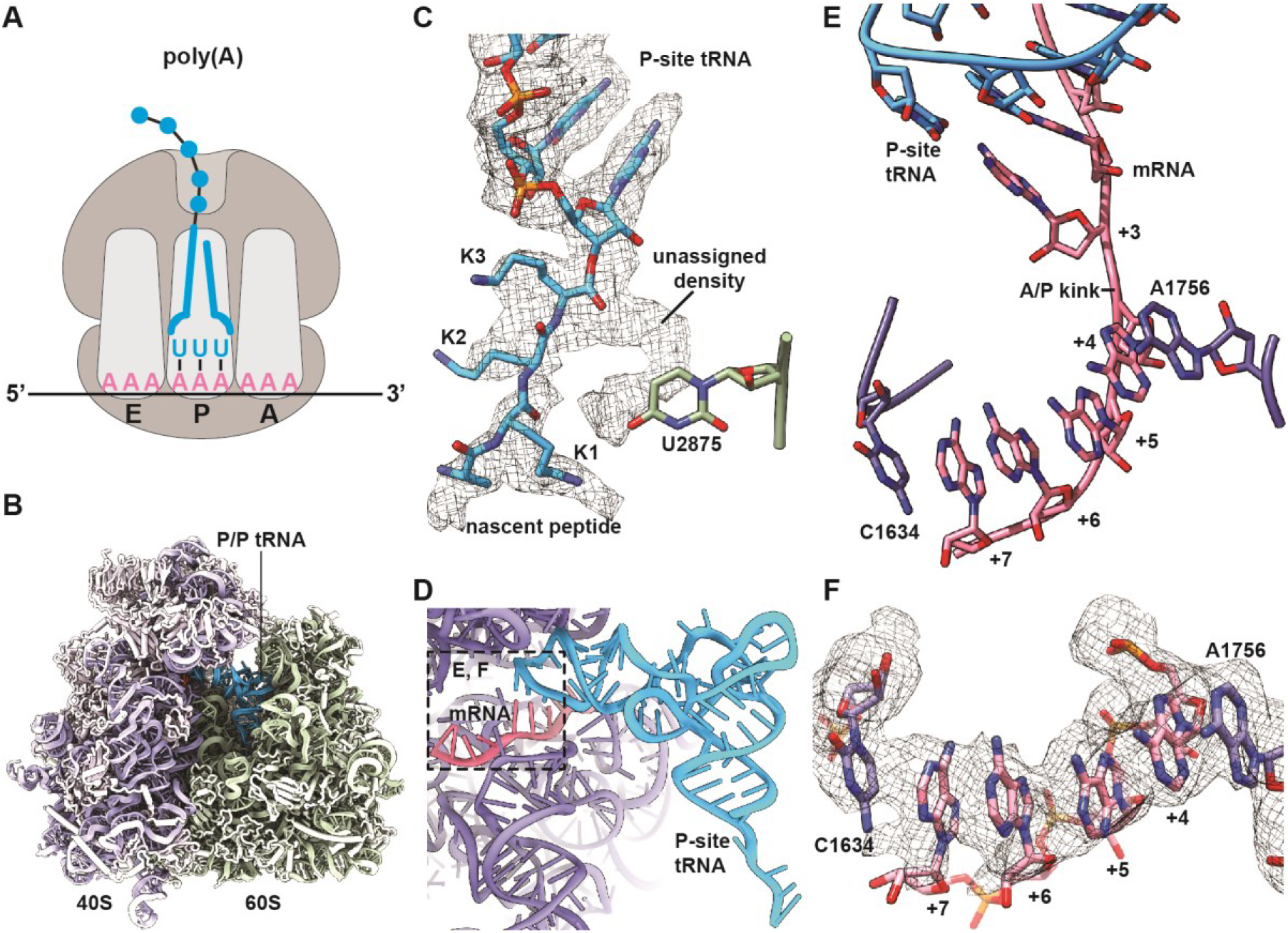
Ribosomes stalled on poly(A) stretches reveal alterations in both the peptidyl-transferase and decoding centers. (**A**) Schematic representation of the stalling situation on poly(A) tract mRNA. Cryo-EM density map of the poly(A) stalled RNC filtered according to local resolution and used to build the molecular model (**B**). (**C**) Cryo-EM density (mesh) and stick model with cartoon phosphate backbone representing the peptidyl-tRNA in the peptidyl-transferase center (PTC). (**D**) General overview of the A, P, and E sites with the P/P tRNA and mRNA. (**E, F**) Detailed view of the mRNA in the A site using sticks model with cartoon phosphate backbone representation and cryo-EM density (mesh). The poly adenine mRNA sequence forms a π-stacking array between positions +4 and +7, which is stabilized from both sides by stacking of 18S rRNA bases C1634 and A1756.

In the resulting structure, we first analyzed the PTC to look for potential structural changes that might rationalize previous arguments that sequential lysines in the peptide tunnel lead to translational stalling due to their basic nature (Lu & Deutsch, 2008). We were able to model the last three C-terminal residues of the nascent chain as lysines, consistent with the RNC being stalled on the poly(A) tract. In the PTC we observed the terminal lysine side chain pointing towards the A site and an extra density not explained by the nascent peptide model (Fig 6C). Overall, however, the crucial catalytic bases (U2875 and U2954) did not seem to be hindered from moving into the induced state conformation upon tRNA binding in the A site, therefore hinting that any perturbations of the PTC geometry are relatively modest. Consistent with this hypothesis, these complexes are reactive to puromycin (data not shown). Moreover, these observations do not provide an explanation for the absence of A site tRNA in 93% of particles. Therefore, we investigated the mRNA conformation in the A site decoding center.

When we examined the molecular details in the decoding center, we clearly saw the structure of the codon-anticodon interaction between the AAA codon and UUU tRNA^Lys^ in the P site with no apparent perturbations (Fig 6D). Strikingly, however, the four downstream adenosines in the A site decoding center are engaged in a π-stacking array, adopting essentially the same single stranded helical conformation recently reported by Passmore and colleagues for isolated poly(A) sequence (Tang, Stowell et al., 2019). This +4 to +7 π-stack is stabilized on both sides by flipped out rRNA nucleotides A1756 and C1634. Indeed, C1634 (C1397 in *E. coli*) is found in an unusual, previously unobserved conformation (Fig 6E and F). In this arrangement, the AAA codon in the A site adopts what is clearly a decoding-incompetent conformation that likely directly contributes to poly(A) mediated stalling, although the general path of mRNA does not seem to be as strongly affected as in the case of both inhibitory codon pairs (Fig EV5C). Taken together, for RNCs stalled on poly(A), we observe structural changes assumed by the mRNA in the A site that preclude canonical interactions with the decoding tRNA.

### Ribosome collisions on poly(A) tracts affect disome formation

Given that ribosome collisions have been shown to produce crucial substrates for quality control pathways (Ikeuchi, Tesina et al., 2019a, Juszkiewicz, Chandrasekaran et al., 2018a, Simms et al., 2017), we wondered if poly(A) tracts in our system would generate a stable ribosome collision amenable to structural analysis. Therefore, we prepared a disome fraction of the poly(A) stalled RNCs as a minimal ribosome collision species and determined structural information by cryo-EM (Appendix Fig S6). We processed the data using the 80S extension approach as described previously (Ikeuchi et al., 2019a) and segregated classes of ribosomal particles stalled in the POST and PRE states (Appendix Fig S9). When we further sorted particles corresponding to the above described poly(A) stalled 80S POST state class, we observed disome structures as expected, however, these POST state ribosomes were found in both the first “stalled” as well as the second “colliding” positions. These collided disomes, which were composed of two POST state ribosomes, are thus strikingly different from previously characterized disomes in both mammalian and yeast systems (Ikeuchi et al., 2019a, Juszkiewicz et al., 2018a). In these previous structures, the second colliding ribosome was always present in a rotated PRE state, with tRNAs in the A/P and P/E states unable to translocate any further downstream. We refined the disome class containing the colliding 80S in the POST state to an overall resolution of 3.8 Å and clearly confirmed that both individual 80S ribosomes are present in the canonical POST state conformation in this disome assembly (Fig 7A–C). Direct comparison of POST-POST with the POST-PRE disome assemblies showed that the second colliding ribosome would have to rotate by 16° to structurally mimic the previously reported POST-PRE conformation (Fig 7D). Taken together, these data indicate that the second colliding ribosome is able to complete the translocation step along the mRNA, a step that would normally be prevented by the stable “roadblock” of the leading stalled ribosome. Therefore, we suggest that poly(A) tracts, which are known to be slippery and allow for sliding, can result in a less rigidly arrested first stalling ribosome.

**Figure 7:**
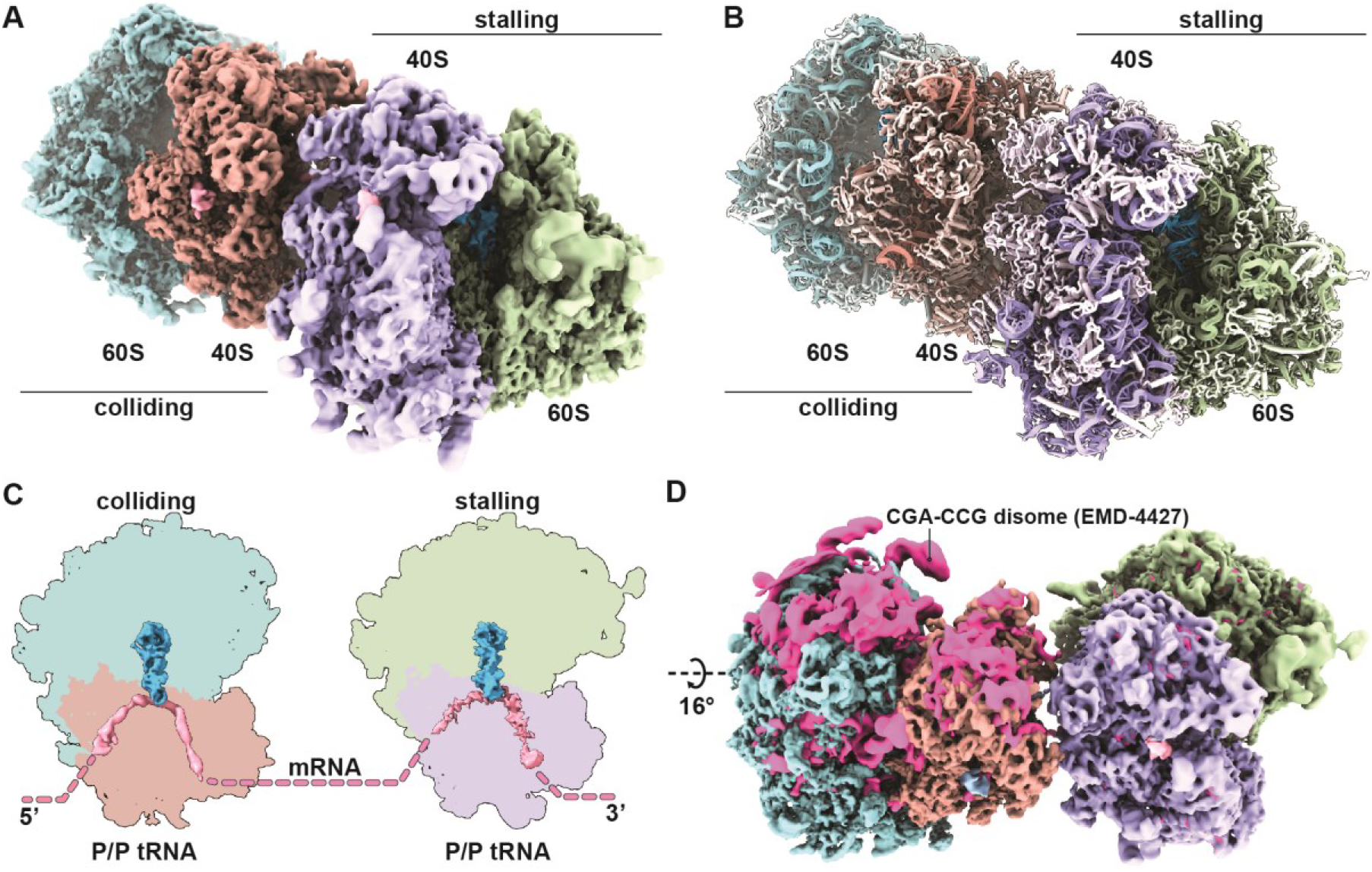
Disomes stalled on poly(A) tracts form a novel POST-POST assembly. (**A**) Composite cryo-EM density map of the POST-POST disome stalled on the poly(A) mRNA reporter filtered according to local resolution and used to build the molecular model (**B**). (**C**) Cut top views of both the first (stalling) and the second (colliding) ribosomes forming the disome. Observed ribosomal and tRNA translocation states are indicated. (**D**) Comparison of ribosomal assemblies between the previously described CGA-CCG stalled yeast disome in pink (EMD-4427, Ikeuchi, Tesina et al., 2019a) and the novel POST-POST assembly observed in poly(A) stalling. The EMD-4427 density map was fitted into the density of the first stalling ribosome on the poly(A) reporter. The indicated rotation was calculated using the 60S subunit, as the compared colliding ribosomes are not in the same translocation state (PRE vs. POST).

## Discussion

Gene expression can be fine-tuned by the selection of specific codons within the context of the degeneracy of the genetic code. While traditional metrics like the codon adaptation index or tRNA adaptation index take into account how commonly a codon is used or how abundant its cognate tRNA is, respectively, it is not well understood why specific codon pairs are underrepresented in genomes compared to their expected values based on the frequency of each individual codon in the pair (Fedorov, Saxonov et al., 2002, Yarus & Folley, 1985). The work of Grayhack and co-workers (Gamble et al., 2016) identified 17 codon pairs in *S. cerevisiae* that reduce protein expression, offering experimental insights into how codon pairs affect translation. In particular, they showed that tRNAs in neighboring ribosomal A and P sites can interact to limit protein output in a codon pair-mediated way, and hypothesized that wobble base pairing played a role in this inhibition.

Our results with an *in vitro* reconstituted translation system directly show that elongation rates of inhibitory codon pairs are slower than those of their optimal counterparts, confirming the hypothesis that inhibition is intrinsic to the ribosome and is likely to involve interactions with the tRNA substrates. For both the Arg-Arg (CGA-CGA) and Arg-Pro (CGA-CCG) pairs, strong defects in the rates and endpoints of the reactions are observed (Figs 2A and B). The observation that the strong endpoint defects are not affected by increased tRNA concentration suggests that there are fundamental structural defects that preclude A site binding/reactivity for some fraction of the ribosome complexes. Consistent with previous work by Grayhack and co-workers (Gamble et al., 2016), the unique I-A wobble associated with decoding CGA codons by the ICG tRNA^Arg^ has a strong effect on interactions in the P site that structurally extend into perturbations of the A site (Figs 5F and I). We additionally find that these defects are partially rescued by substitution of a UCG tRNA^Arg^ that no longer relies on I:A pairing (Fig 2C), consistent with previous *in vivo* studies (Gamble et al., 2016).

The observation that the kinetics of decoding are retarded by inhibitory codon pairs in biochemical assays was corroborated by our high resolution ribosome profiling studies. We see an enrichment of 21 nt RPFs, corresponding to ribosomes lacking a tRNA in the A site, when the first codon of the pair is in the ribosomal P site and the second codon is in the A site (Fig 3A and B). Comparing the results from the CHX/ANS library with the CHX/TIG library, we see the same level of these 21 nt RPFs, indicating that peptide bond formation is not limiting the inhibitory codon pair-stalled ribosomes (Fig 3C). This observation is consistent with the fact that the optimal codon pairs (which encode the same amino acid residues and use the same tRNAs) elongate at normal rates both *in vivo* and *in vitro*. These data indicate that the inhibitory codon pairs affect the decoding center of the 40S subunit rather than the peptidyl-transferase center of the 60S subunit. Overall, our data are consistent with the idea that the major mechanism of inhibition on most of these inhibitory codon pairs is through impairment of tRNA binding/accommodation.

Previous studies argue that ribosomes lacking the ribosomal protein Asc1 are able to readthrough CGA-CGA codons, thus effectively increasing protein output (Letzring et al., 2013, Wang et al., 2018, Wolf & Grayhack, 2015). Our data argue that this apparent gain of function may originate in part from defects in the biochemical activity of ribosomes lacking Asc1. First, we find that ribosomes lacking Asc1 are less efficient at elongating on mRNAs with both inhibitory and optimal pairs *in vitro* (Fig 4A and EV3B). Second, by ribosome profiling, we observe a higher fraction of 21 nt RPFs in cells lacking Asc1 suggesting a general defect in tRNA decoding within the A site of the ribosome (Fig 4B and C). While this finding is somewhat surprising from a structural perspective, given that Asc1 is located on the 40S subunit far away from the decoding center, one possibility is that the loss of Asc1 affects the conformation of Rps3, a ribosomal protein that directly interacts with Asc1 and forms a part of the mRNA entry channel (Limoncelli, Merrikh et al., 2017, Simms, Kim et al., 2018). Asc1 is also positioned such that it may be involved in sensing ribosome collisions that lead to ribosome rescue pathways (Ikeuchi & Inada, 2016, Ikeuchi et al., 2019a, Juszkiewicz et al., 2018a). It seems likely that the increased read-through on inhibitory sequences in the Asc1 deletion strain arises from initial defects in the decoding step (promoting frameshifting) as well as by the loss of cellular responses to ribosome pausing.

Detailed mechanistic insight into the origins of A site accommodation defects was ultimately provided by our structural analysis. In our cryo-EM structures of ribosome-nascent chain complexes stalled on the CGA-CGA or CGA-CCG codon pairs, we identified several structural details that likely directly affect tRNA binding/accommodation activity. Interestingly, in each case these alterations are mediated by the structure of the mRNA itself and readily explain the previously proposed communication between the ribosomal A- and P sites (Figs 5 and EV5) (Gamble et al., 2016). In particular, for both the CGA-CCG and CGA-CGA inhibitory pairs, the C+4 mRNA nucleotide is dramatically flipped away from the A site decoding center of the ribosome. The C+4 nucleotide instead makes contact with the P site codon and interacts with C1637 of 18S rRNA which stacks to the anticodon inosine decoding the wobble position (A+3) of mRNA in the P site. The path of the mRNA is also affected by the purine:purine A:I wobble base pair at position +3 and shifts towards C1637. This perturbation involving the A:I wobble interaction provides an immediate explanation for why the CGA codon in particular confers the strongest elongation defect. Moreover, the A/P kink of the mRNA, which was shown to be crucial for A site interaction and proofreading (Keedy et al., 2018), is moved downstream in these structures as a consequence (Fig 5F and I). This critical structure is typically stabilized by an ammonium ion in X-ray structures (Rozov, Khusainov et al., 2019) and was proposed to be essential for frame maintenance by preventing slippage (Selmer, Dunham et al., 2006). Finally, in the case of CGA-CCG, we observe a hairpin structure formed by mRNA nucleotides between positions +5 and +14 (Fig EV5D and E). This structure may be particular to this reporter mRNA sequence since no equivalent stable mRNA secondary structure is formed in the case of the CGA-CGA stalled RNC. Interestingly, a similar A site hairpin was observed previously in a structure implicated in translational bypassing (Agirrezabala, Samatova et al., 2017).

Consistent with the earlier work (Gamble et al., 2016), we see a specific deleterious effect of I:A wobble decoding on translation efficiency in inhibitory codon pairs containing the 5’ CGA codon. Previously, the purine:purine I:A base pair was analyzed in the A site only, where its accommodation affects and alters mainly the anticodon of tRNA, due to its unique “wide” purine-purine geometry (Murphy & Ramakrishnan, 2004). In contrast, in our structure of the I:A wobble pair in the P site, we find that its accommodation affects not the anticodon of tRNA but rather the mRNA backbone (Figs 5E and H). This mRNA distortion apparently imposes allosteric effects on the neighboring region resulting in the unusual mRNA conformation in the A site. The modification of adenosine to inosine (Gerber & Keller, 1999) expands the decoding range of the ICG tRNA^Arg^ as inosine is able to base-pair with cytidine, uridine and even adenosine at the wobble position. It is intriguing to observe that this seemingly elegant evolutionary decoding mechanism has certain associated disadvantages as the non-optimal CGA codon (decoded via the I:A interaction with the ICG tRNA^Arg^) is slow to decode and leads to deleterious effects on mRNA stability (Presnyak, Alhusaini et al., 2015b).

In the case of translation of poly(A) tracts, previous studies proposed that electrostatic interactions between the poly-basic nascent chain and the peptide exit tunnel of the ribosome might elicit ribosomal stalling (Lu & Deutsch, 2008). Using our detailed structural information, we were able to reveal that an mRNA-mediated mechanism is directly contributing to stalling. Consecutive adenosines are engaged in a π-stacking array in the A site, stabilized on both sides by rRNA base stacking interactions, and adopt a helical conformation typical for single stranded poly(A) stretches (Fig 6E and F) (Tang et al., 2019). This π-stacking array represents a decoding-incompetent structure. Conversely, the crucial catalytic bases in the peptidyl-transferase center (PTC) did not seem to be hindered from moving into the induced state conformation despite the presence of extra density which is not clearly interpretable (Fig 6C). This extra density adopts a defined shape next to the last nascent amino acid residue and could potentially be assigned to a mixed nascent chain state or even a small molecule. However, the observed geometry of the PTC cannot explain the highly efficient stalling on poly(A) tracts and the absence of any A site tRNA in 93% of particles in the dataset. Therefore, we argue that the inhibitory conformation of mRNA in the A site is at the basis of the poly(A)-mediated stalling mechanism. These ideas agree with previous observations that consecutive AAG codons are less efficient in stalling than AAA codons (Koutmou et al., 2015) despite encoding for the same amino acid residue and that the intrinsic π-stacked helical structure of poly(A) single strand tract is efficiently disrupted by inclusion of guanosines (Tang et al., 2019). Taken together, while we can’t exclude the possibility that the basic nascent chain also contributes, the stalling mechanism employed at poly(A) stretches mainly depends on the specific inhibitory conformation of the mRNA in the A site.

Interestingly, when studying ribosomal collisions as a consequence of poly(A)-mediated stalling, we found a large fraction of the disomes in a novel POST-POST state that was distinct from the previously characterized disome structures in both mammalian and yeast systems (Fig 7A and B) (Ikeuchi et al., 2019a, Juszkiewicz et al., 2018a). In both previous structures, the second colliding ribosome is captured in a rotated PRE state unable to translocate further. Finding both collided ribosomes in the POST state indicates that the second colliding ribosome completed the translocation step, likely due to a weaker “roadblock” presented by the first stalled ribosome. ince poly(A) tracts were characterized as slippery (Koutmou et al., 2015), it is tempting to speculate that applying force on the first stalled ribosome by the colliding ribosome(s) could contribute to ribosome sliding on the mRNA and loss of reading frame. This model is consistent with recent findings that directly implicate ribosomal collisions in +1 frameshifting (Simms, Yan et al., 2019). Ribosomal collisions could, in principle, disrupt the interaction between the P site tRNA and the mRNA in the first ribosome and contribute to +1 frameshifting observed after ribosomal pausing (Dinman, 2012). We speculate that the loss of reading frame in the case of collisions on poly(A) tracts is facilitated by (i) the fact that the P site tRNA is the only one left on the stalled ribosome after the E site tRNA dissociates and (ii) the fact that the P site tRNA only interacts with the mRNA via relatively less stable A:U base pairs. These ideas are consistent with earlier studies arguing that reading frame maintenance is predominantly affected by the energetics of the P-site codon-anticodon interaction (Baranov, Gesteland et al., 2004).

Taken together, our work combines *in vitro* and *in vivo* methods to study the effects of inhibitory mRNA sequences, and shows for the first time detailed mechanistic insight into mRNA-mediated translation stalling via decoding obstruction.

## Methods

### Ribosome Preparation

WT Ribosomes were purified and isolated as subunits as previously described (Eyler & Green, 2011). Asc1 depleted ribosomes were purified similarly from strain AW768 (MATa his3-Δ1, leu2-Δ0, met15-Δ0, ura3-Δ0, asc1-Δ::spHIS5, pURA3, ASC1) gifted from the Grayhack lab (Wolf & Grayhack, 2015).

### Purification of translation factors

Translation initiation factors eIF1, eIF1A, eIF5, eIF5B were expressed and purified from *E. coli* and eIF2 was expressed and purified from S. cerevisiae as previously described (Acker, Kolitz et al., 2007, Eyler & Green, 2011). The translation elongation factor, eIF5A was purified from *E. coli* as previously described (Gutierrez, Shin et al., 2013, Schuller et al., 2017). The translation elongation factors eEF2 and eEF3 were purified from S. cerevisiae as previously described (Schuller et al., 2017).

### Purification of amino-acyl synthetases

Plasmids gifted from the Grayhack lab containing the arginine and proline sythetases were transformed into BY4741 yeast strain and grown initially in CSM –ura glucose media (Sunrise Science) and induced in –ura galactose media overnight. Harvested cells grown in small scale (500 mL) were lysed by vortexing with acid washed glass beads (sigma) in extraction buffer (50mM Tris-Cl, pH 7.5, 1M NaCl, 1mM EDTA, 4mM MgCl2, 5mM DTT, 10% glycerol). Larger scale preparations (2 L) were lysed by CryoMill and lysate was flowed over 5mL Ni column (GE) and batch eluted in 5 to 10 mLs (extraction buffer used for lysis with 5mM BME rather than DTT). Lysates were then diluted in IPP0 buffer (10mM Tris-Cl, pH 8, 0.1% NP40) and incubated for a minimum of 2 hours with IgG sepharose beads at 4°C. Beads were spun down at low speed (2 krpm) and unbound supernatant was removed. The beads were then washed with multiple times with IPP150 buffer (10 mM Tris-Cl, pH 8, 150mM NaCl, 0.1% NP40) to remove all unbound protein and washed subsequently with cleavage buffer (10 mM Tris-Cl, pH 8, 150 mM NaCl, 0.1% NP40, 0.5mM EDTA, 1 mM DTT). The protein was then cleaved from the beads using 3C protease in cleavage buffer overnight at 4°C. Cleaved protein was removed from beads, flash froze in small aliquots and stored at −80°C for use.

### Purification of bulk yeast tRNA

tRNA isolation protocol was derived from a protocol to isolate RNA from *E. coli* (Ehrenstein, 1967) with minor changes and an added LiCl precipitation to remove rRNA and mRNA. Briefly, 3L of BY4741 yeast alone or expressing a plasmid of interest were grown to an OD600 of 1 and harvested by centrifugation. Cell pellets were resuspended in 20 mL Buffer A (50mM NaOAc, pH 7.5, 10mM MgOAc). Phenol:chloroform extraction of RNA and DNA was performed using an equal volume of acid phenol:chloroform, pH 4.5 (VWR). rRNA and mRNA was then pelleted by LiCl precipitation and tRNA and DNA was then ethanol precipitated. DNA was then removed by isopropyl alcohol precipitation. tRNA was then deacylated by incubation in 1M Tris-Cl, pH 9 for 3 hours at room temperature. Deacylated tRNA was then purified by ethanol precipitation and resuspended in water for acylation and use in *in vitro* assays.

### Purification and charging of tRNAs

Initiator methionine and lysine tRNAs were purchased from tRNA probes (College Station, TX). Phenylalanine tRNA was purchased from Sigma. Arginine and proline tRNAs were isolated from bulk yeast tRNA using 3’ biotinylated oligonucleotides (listed below) as previously described (Yokogawa et al., 2010).

Oligo for A(I)CG-tRNA^Arg^: 5’ – CGC AGC CAG ACG CCG TGA CCA TTG GGC – 3’ Biotin

Oligo for UGG-tRNA^Pro^: 5’ – CCA AAG CGA GAA TCA TAC CAC TAG AC – 3’ Biotin

Leu-2um plasmids for overexpressing native and exact match tRNAs were received from the Grayhack lab (ECB0873 ACG-tRNA^Arg^, ECB0874 UCG-tRNA^Arg^). tRNA sequences were moved to pRS316 vector by Gibson cloning for lower level overexpression. The low copy CEN plasmids containing the tRNA sequences were transformed into the BY4741 yeast strain. Bulk tRNA was then purified by the protocol above and the non-native tRNA was then isolated by the same 3’ biotinylated oligonucleotide method previously (Yokogawa et al., 2010) using the specific oligonucleotides listed below.

Oligo for A(I)CG-tRNA^Arg^: 5’ – CGC AGC CAG ACG CCG TGA CCA TTG GGC – 3’ Biotin

Oligo for UCG-tRNA^Arg^: 5’ – CGA AGC CAG ACG CCG TGA CCA TTG GGC – 3’ Biotin

All isolated tRNAs were subjected to CCA addition as described previously (Gutierrez et al., 2013). Isolated tRNA^Lys^ was charged using S100 extract and tRNA^Phe^, tRNA^Arg^, and tRNA^Pro^ were charged using purified synthetases as previously described with minor changes (Eyler & Green, 2011). Briefly, reactions contained 1X buffer 517 (30 mM HEPES-KOH pH 7.4, 30 mM KCl, 15 mM MgCl_2_), 4 mM ATP, 5 mM DTT, 10-20 µM amino acid, 3 µM CCA-added tRNA and a 1/5 th volume of an S100 extract or 10 µM tRNA synthetase. Reactions were incubated at 30°C for 30 minutes, then extracted twice with acid phenol and once with chloroform. tRNA was precipitated with ethanol, resuspended in 20 mM KOAc, 2 mM DTT, pH 5.2, and stored in small aliquots at −80°C.

### *In vitro* 80S initiation complex formation

80S initiation complexes were formed as previously described (Schuller et al., 2017) with minor differences. Briefly, 3 pmol of 35S-Met-tRNAiMet was mixed with 50 pmol of eIF2 and 1 mM GTP in 1X Buffer E (20 mM Tris pH 7.5, 100 mM KOAc pH 7.6, 2.5 mM Mg(OAc)_2_, 0.25 mM Spermidine, and 2 mM DTT) for 10 min at 26°C. Next a mixture containing 25 pmol 40S subunits, 200 pmol mRNA (purchased from IDT), 125 pmol eIF1, and 125 pmol eIF1A in 1X Buffer E was added for 5 min. To form the 80S complex, a mixture containing 25 pmol 60S subunits, 150 pmol eIF5, 125 pmol eIF5b, and 1 mM GTP in 1X Buffer E was added for 1 min. Complexes were then mixed 1:1 with buffer E containing 17.5 mM Mg(OAc)_2_ to yield a final magnesium concentration of 10 mM. Ribosomes were then pelleted through a 600 μL sucrose cushion containing 1.1 M sucrose in buffer E with 10 mM Mg(OAc)_2_ using a MLA-130 rotor (Beckmann) at 75,000 rpm for 1 hr at 4°C. After pelleting, ribosomes were resuspended in 15-25 μL of 1X Buffer E containing 10 mM Mg(OAc)_2_ and stored at −80°C.

### *In vitro* reconstituted translation elongation

Translation elongation reactions were performed as previously described (Eyler & Green, 2011, Schuller et al., 2017) with minor differences. Briefly, aa-tRNA ternary complex was formed by incubating aa-tRNA (1.5 – 2 uM), eEF1A (5 uM), 1 mM GTP, in 1X Buffer E for 10 minutes at 26°C. Limited amounts of 80S initiation complexes (3 nM) were then mixed with aa-tRNA ternary complex (varying concentrations), eEF2 (500 nM), eEF3 (1 μM), eIF5A (1 μM), ATP (3 mM) and GTP (2 mM). Reactions were incubated at 26°C and time points quenched into 500mM KOH. Samples were diluted 1 uL into 3 uL water before monitoring peptide formation electrophoretic TLC (Millipore). TLC plates were equilibrated with pyridine acetate buffer (5 mL pyridine, 200 mL acetic acid in 1 L, pH 2.8) before electrophoresis at 1200 V for 25 to 30 minutes. Plates were developed using a Typhoon FLA 9500 Phosphorimager system (GE Healthcare Life Sciences) and quantified using ImageQuantTL (GE Healthcare Life Sciences). Time courses were fit to single exponential kinetics using Kaleidagraph (Synergy Software).

### *In vitro* Met-Puromycin assay

Reactions were set up as previously described (Schuller et al., 2017). Reactions were performed for each set of initiation complexes made and used to normalize peptide formation from elongation. Briefly, 2 nM initiation complexes and 1 μM eIF5A in 1X Buffer E (20 mM Tris pH 7.5, 100 mM KOAc pH 7.6, 2.5 mM Mg(OAc)_2_, 0.25 mM Spermidine, and 2 mM DTT) were incubated at 26°C in the presence of 4 mM puromycin. Time points over the course of 120 min were quenched into 500 mM KOH and analyzed by electrophoretic TLC (Millipore). TLC plates were equilibrated with pyridine acetate buffer (5 mL pyridine, 200 mL acetic acid in 1 L, pH 2.8) before electrophoresis at 1200 V for 25 min. Plates were developed using a Typhoon FLA 9500 Phosphorimager system (GE Healthcare Life Sciences) and quantified using ImageQuantTL (GE Healthcare Life Sciences).

### *In vitro* PTH assay to access peptidyl-tRNA drop-off

Translation elongation reactions were performed in the presence of 27 μM peptidyl-tRNA hydrolase (PTH) to monitor drop-off of peptidyl-tRNAs from translating ribosomes as described previously (Schuller et al., 2017). Time points for drop-off products were quenched with 10% formic acid and were analyzed by electrophoretic TLC in pyridine acetate buffer (see above) at 1200 V for 30 minutes.

### Preparation of ribosome footprint libraries and analysis of aligned footprints

WT and ∆asc1 cells were grown to OD ~ 0.5 in 1 L of YPD media (sample 1) or transferred to YPGR media (2% galactose and 2% raffinose) for 6 hr (sample 2) and harvested by fast filtration followed by flash frozen in liquid nitrogen. Cell pellets were ground with 1 mL footprint lysis buffer [20 mM Tris-Cl (pH8.0), 140 mM KCl, 1.5 mM MgCl_2_, 1% Triton X-100 0.1 mg/mL CHX, 0.1 mg/mL TIG] in a Spex 6870 freezer mill. Lyzed cell pellets were diluted to 15 mL in footprint lysis buffer and clarified by centrifugation. Polysomes were isolated from sucrose cushions for library construction as described previously (Wu et al., 2019).

3’ adapter (NNNNNNCACTCGGGCACCAAGGA) was trimmed, and 4 random nucleotides included in RT primer were removed from the 5’ end of reads (RNNNAGATCGGAAGAGCGTCGTGTAGGGAAAGAGTG TAGATCTCGGTGGTCGC/iSP18/TTCAGACGTGTGCTCTTCCGATCTGTCCTTGGTGCCCGAGTG). Trimmed reads longer were aligned to yeast ribosomal and non-coding RNA sequence. Unmapped reads were mapped to R64-1-1 S288C reference genome assembly (SacCer3) from the *Saccharomyces* Genome Database Project using STAR (Dobin, Davis et al., 2013) as described previously (Wu et al., 2019). Data shown in Figs 3 and 4 for WT are identical to those published previously (Wu et al., 2019). Relative ribosome occupancies for codon pairs were computed by taking the ratio of the ribosome density in a 3-nt window at the di-codon over the density in the coding sequence (excluding the first and the last 15 nt).

### Preparation of stalled ribosome-nascent chain complexes

We generated a series of mRNA reporters containing three different stalling sequences (CGA-CCG)_2_, (CGA-CGA)_2_, and poly(A) (Appendix Figs S1A, S2A and S6A). These sequences were placed downstream of a sequence coding for TEV-cleavable N-terminal His- and HA tags and the first 64 amino acid residues of truncated uL4. Corresponding mRNAs were produced using the mMessage mMachine Kit (Thermo Fischer) utilizing an upstream T7 promoter and translated in a yeast cell-free translation extract from *ski2*Δ cells.

This yeast translation extract was prepared, and *in vitro* translation was performed essentially as described before (Waters & Blobel, 1986). In brief, the cells were grown in YPD medium to OD_600_ of 1.5–2.0. Spheroplasts were prepared from harvested washed cells using 10 mM DTT for 15 min at room temperature and 2.08 mg zymolyase per 1 g of cell pellet for 75 min in 1 M sorbitol at 30°C. Spheroplasts were then washed and lysed in a Dounce homogenizer as described (Waters & Blobel, 1986) before using lysis buffer comprising 20 mM Hepes pH 7.5, 100 mM KOAc, 2 mM Mg(OAc)_2_, 10% glycerol, 1 mM DTT, 0.5 mM PMSF and complete EDTA-free protease inhibitors (GE Healthcare). The S100 fraction of lysate supernatant was passed through PD10 column (GE Healthcare) and used for *in vitro* translation. *In vitro* translation was performed at 17°C for 75 min using great excess of template mRNA (38 µg per 415 µl of extract) to prevent degradation of resulting stalled ribosomes by endogenous response factors.

Respective stalled RNCs were affinity-purified using the His_6_^−^ tag of the nascent polypeptide chain essentially as described before (Ikeuchi et al., 2019a, Tesina, Heckel et al., 2019). After *in vitro* translation, the extract was applied to Ni-NTA Dynabeads^TM^ (Invitrogen) and incubated while rotating for 15 min at 4°C. The beads were washed three times with excess of a wash buffer containing 50 mM HEPES/KOH, pH 7.5, 100 mM KOAc, 25 mM Mg (OAc)_2_, 250 mM sucrose, 0.1% Nikkol and 5 mM ß-Mercaptoethanol and eluted in 400 µl of the same buffer containing 300 mM imidazole. The elution was applied to a 10-50% sucrose gradient in wash buffer, and ribosomal fractions were separated by centrifugation for 3 h at 172,000 g at 4°C in a SW40 rotor. For gradient fractionation, a Piston Gradient Fractionator^TM^ (BIOCOMP) was used. The 80S (mono)ribosome (and for poly(A) also the disome) fractions were collected, applied onto 400 µl of sucrose cushion buffer and spun at 534,000 g for 45 min at 4°C in a TLA110 rotor. The resulting ribosomal pellets were resuspended carefully on ice in 25 µl of grid buffer (20 mM HEPES/KOH, pH 7.2, 50 mM KOAc, 5 mM Mg(OAc)_2_, 125 mM sucrose, 0.05% Nikkol, 1 mM DTT and 0.01 U/µl SUPERase-IN^TM^ (Invitrogen).

Collected 80S fractions of CGA-CCG and CGA-CGA stalled RNCs were also subjected to puromycin reactions with 1 mM puromycin at 20°C. Time point samples were heated 5 minutes at 60°C with reducing sample buffer and analyzed by SDS-PAGE and western blotting.

### Electrophoresis and Western blotting

Protein samples of *in vitro* translation reactions and subsequent purifications were separated on SDS-PAGE at neutral pH condition (pH 6.8, for purified protein samples) and were transferred on PVDF membrane (Immobilon-P, Millipore). After blocking with 5% skim milk in PBS-T, the membranes were incubated with anti-HA-peroxidase antibody (1:5,000; Roche, Cat# 12013819001, clone 3F10) for 1 h at room temperature followed by washing with PBS-T for three times. Chemiluminescence was detected using SuperSignal® substrate (Thermo Fischer) in a LAS4000 mini (GE Healthcare).

### Cryo-EM

Freshly prepared samples (stalled monosomes or disomes) were applied to 2 nm pre-coated Quantinfoil R3/3 holey carbon support grids and vitrified. Data were collected at Titan Krios TEM (Thermo Fisher) equipped with a Falcon II direct detector at 300 keV under low-dose conditions of about 25 e-/Å2 for 10 frames in total and defocus range of −1.3 to −2.8 µm. Magnification settings resulted in a pixel size of 1.084 Å/pixel. In the case of CGA-CGA RNCs, a higher magnification was used resulting in a pixel size of 0.847 Å/pixel. Original image stacks were summed and corrected for drift and beam-induced motion at the micrograph level by using MotionCor2 (Zheng, Palovcak et al., 2017). The Contrast transfer function (CTF) estimation and resolution range of each micrograph were performed with Gctf (Zhang, 2016).

### Cryo-EM Data processing

All datasets were processed using standard procedures with programs GAUTOMATCH (http://www.mrc-lmb.cam.ac.uk/kzhang/) used for particle picking and RELION-3 for data processing and 3D reconstruction (Zivanov, Nakane et al., 2018). For each dataset, picked particles were extracted for 2D classification using a box of 400 pixels rescaled to 70 pixels. After selection of suitable 2D classes, particles were extracted for initial 3D refinement followed by 3D classification using a box of 400 pixel rescaled to 120 pixels and a mask diameter of 300 Å.

The CGA-CCG dataset was described before with focus on the Xrn1 factor bound (Tesina et al., 2019). We now re-processed this dataset with focus on the ribosome itself. Individual translation states were separated as before with around 60% of the particles containing tRNAs in the P/P and E/E conformation (Appendix Fig S3). These classes were joined and separated into four subclasses sorting out low resolution particles. Further subclassification was performed using a mask covering tRNAs. This approach sorted out a population of particles without the E site tRNA. The cleaned population of particles was further processed using particle CTF refinement yielding a final resolution of 2.6 Å. This cryo-EM density map was filtered according to local resolution and used for interpretation (Appendix Fig S8A).

For the CGA-CGA dataset, 840,234 particles were used after 2D classification and sorted into six classes in 3D classification. A vast majority of programmed ribosomal particles in the dataset were found in the post translocation state while a single class containing tRNAs in P/P E/E conformation represented 39.9% of the whole dataset (Appendix Fig S4). As further classification of this class was mainly yielding volumes sorted based on position of expansion segment 27 on the periphery of the ribosome, the class was further processed as a whole. The final cryo-EM density map reaching an overall resolution of 3.2 Å after particle CTF refinement was filtered according to local resolution and used for interpretation (Appendix Fig S8B).

For the poly(A) 80S dataset, 840,234 particles were used after 2D classification and sorted into six classes in 3D classification (Appendix Fig S7). Analogous to previous datasets, a vast majority of programmed ribosomal particles represented classes in the post translocation state. Class 3 containing tRNAs in P/P E/E conformation was subsorted based on tRNA presence into classes containing only P/P tRNA and a class containing both P/P and E/E tRNAs. The dominant classes of P/P tRNA state were joined and further processed using particle CTF refinement and Bayesian polishing. The resulting cryo-EM density map reached an overall resolution of 3.1 Å. This volume was subjected to focused refinement using a mask covering the 60S subunit and the decoding center. This yielded a better resolved density map (3.0 Å) in the region of interest and was used for interpretation after filtering according to local resolution (Appendix Fig S8C).

### Reconstruction of the poly(A) disome

The poly(A) disome dataset was collected as described above. The dataset was processed using the “80S extension” approach as described previously (Ikeuchi et al., 2019a). Initial 3D classification yielded in a class of ribosomes in the POST state with P/P tRNA as described above for the poly(A) monosome. Surprisingly, subsorting of this class revealed that approximately the same share of particles in this class represented the first stalling and the second colliding ribosome judging by the density of the neighboring ribosome close to the mRNA entry and exit site, respectively (Appendix Fig S9). Further processing of the leading POST state ribosome (with neighbor density at mRNA exit) yielded a standard POST-PRE hybrid disome assembly as observed for the CGA-CCG-stalled disome (Ikeuchi et al., 2019a). On the other hand, processing of the second colliding ribosome in the POST state (with neighbor density at mRNA entry) revealed a novel POST-POST disome assembly. Both these volumes were obtained by stepwise box extension and refinement with particle re-centering (fist 500 pixels rescaled to 120 pixels followed by 700 pixels rescaled to 506 pixels). Soft masks covering individual ribosomal bodies were used for multi-body refinement to obtain a more detailed information (Nakane, Kimanius et al., 2018). The resulting volumes were filtered according to local resolution (Appendix Fig S10) and fitted into the consensus refinement yielding a composite cryo-EM density map at 3.8 Å overall resolution.

### Model building

To generate molecular models for our structures, we used our previously refined models for stalled yeast ribosomes (Tesina et al., 2019) PDB ID: 6Q8Y and (Ikeuchi et al., 2019a) PDB ID: 6I7O). First, individual subunits and tRNAs were fitted as rigid bodies into the densities. These models were then refined and remodeled in COOT (Brown, Long et al., 2015) and Phenix (Adams, Afonine et al., 2010). Cryo-EM structures and models were displayed with UCSF Chimera (Pettersen, Goddard et al., 2004) and ChimeraX (Goddard, Huang et al., 2018). Detailed statistics of model refinements and validations are listed in Appendix Tables 1-X.

## Supporting information

Expanded view figures

## Data availability

The cryo-EM structures reported here have been deposited in the Protein Data Bank under accession codes XXXXXX and in the Electron Microscopy Data Bank under accession codes YYYYYY. Ribosome profiling datasets have been deposited under GSE136202 (reviewer access with secure token: mfwfweqatvmlfkt).

## Acknowledgements

The authors would like to thank Elisabeth Heckel for help with sample preparation and to Charlotte Ungewickell and Susanne Rieder for technical support. This study was supported by a grant of the Deutsche Forschungsgemeinschaft (DFG; BE1814/15-1) to RBe and a Ph.D. fellowship from Boehringer Ingelheim Fonds to RBu.

## Author contributions

RG, RBe, TB and AB conceived the study and designed experiments. Experiments were performed by LL, PT and CW. LL performed yeast *in vitro* kinetics experiments and analyzed the corresponding data. CW performed experiments and data analysis for ribosome profiling. PT prepared cryo-EM samples and processed the data, OB collected the data. RBu built atomic models and JC, PT and TB refined them. PT, LL, AB, TB, RBe and RG wrote the manuscript.

## References

Acker MG, Kolitz SE, Mitchell SF, Nanda JS, Lorsch JR (2007) Reconstitution of yeast translation initiation. Methods in enzymology 430: 111–145

Adams PD, Afonine PV, Bunkoczi G, Chen VB, Davis IW, Echols N, Headd JJ, Hung LW, Kapral GJ, Grosse-Kunstleve RW, McCoy AJ, Moriarty NW, Oeffner R, Read RJ, Richardson DC, Richardson JS, Terwilliger TC, Zwart PH (2010) PHENIX:a comprehensive Python-based system for macromolecular structure solution. Acta crystallographica Section D, Biological crystallography 66: 213–21

Agirrezabala X, Samatova E, Klimova M, Zamora M, Gil-Carton D, Rodnina MV, Valle M (2017) Ribosome rearrangements at the onset of translational bypassing. Sci Adv 3: e1700147

Arthur L, Pavlovic-Djuranovic S, Smith-Koutmou K, Green R, Szczesny P, Djuranovic S (2015) Translational control by lysine-encoding A-rich sequences. Sci Adv 1

Arthur LL, Djuranovic S (2018) PolyA tracks, polybasic peptides, poly-translational hurdles. Wiley Interdiscip Rev RNA: e1486

Baranov PV, Gesteland RF, Atkins JF (2004) P-site tRNA is a crucial initiator of ribosomal frameshifting. RNA 10: 221–30

Behrmann E, Loerke J, Budkevich TV, Yamamoto K, Schmidt A, Penczek PA, Vos MR, Bürger J, Mielke T, Scheerer P, Spahn CM (2015) Structural snapshots of actively translating human ribosomes. Cell 161: 845–857

Brandman O, Hegde RS (2016) Ribosome-associated protein quality control. Nature structural & molecular biology 23: 7–15

Brown A, Long F, Nicholls RA, Toots J, Emsley P, Murshudov G (2015) Tools for macromolecular model building and refinement into electron cryo-microscopy reconstructions. Acta crystallographica Section D, Biological crystallography 71: 136–53

Brule CE, Grayhack EJ (2017) Synonymous Codons: Choose Wisely for Expression. Trends in genetics: TIG 33: 283–297

Burgess-Brown NA, Sharma S, Sobott F, Loenarz C, Oppermann U, Gileadi O (2008) Codon optimization can improve expression of human genes in Escherichia coli: A multi-gene study. Protein expression and purification 59: 94–102

Buschauer A, Matsuo Y, Chen Y, Alhusaini N, Sweet T, Sugiyama T, Ikeuchi K, Cheng J, Matsuki Y, Gilmozzi 857 A, Berninghausen O, Becker T, Coller J, Inada T, Beckmann R (2019) The Ccr4-Not complex monitors the 858 translating ribosome for codon optimality. Manuscript under revision, Science

Dana A, Tuller T (2014) The effect of tRNA levels on decoding times of mRNA codons. Nucleic acids research 42: 9171–9181

Dinman JD (2012) Mechanisms and implications of programmed translational frameshifting. Wiley Interdiscip Rev RNA 3: 661–73

Dobin A, Davis CA, Schlesinger F, Drenkow J, Zaleski C, Jha S, Batut P, Chaisson M, Gingeras TR (2013) STAR: ultrafast universal RNA-seq aligner. Bioinformatics (Oxford, England) 29: 15–21

dos Reis M, Savva R, Wernisch L (2004) Solving the riddle of codon usage preferences: a test for translational selection. Nucleic acids research 32: 5036–5044

Ehrenstein G (1967) [76] Isolation of sRNA from intact Escherichia coli cells. Elsevier,

Elf J, Nilsson D, Tenson T, Ehrenberg M (2003) Selective charging of tRNA isoacceptors explains patterns of codon usage. Science (New York, NY) 300: 1718–1722

Eyler DE, Green R (2011) Distinct response of yeast ribosomes to a miscoding event during translation. RNA 17: 925–932

Fedorov A, Saxonov S, Gilbert W (2002) Regularities of context-dependent codon bias in eukaryotic genes. Nucleic acids research 30: 1192–1197

Frischmeyer PA, van Hoof A, O’Donnell K, Guerrerio AL, Parker R, Dietz HC (2002) An mRNA surveillance mechanism that eliminates transcripts lacking termination codons. Science 295: 2258–61

Gamble CE, Brule CE, Dean KM, Fields S, Grayhack EJ (2016) Adjacent Codons Act in Concert to Modulate Translation Efficiency in Yeast. Cell 166: 679–690

Gerber AP, Keller W (1999) An adenosine deaminase that generates inosine at the wobble position of tRNAs. Science 286: 1146–9

Gingold H, Pilpel Y (2011) Determinants of translation efficiency and accuracy. Molecular systems biology 7: 481

Goddard TD, Huang CC, Meng EC, Pettersen EF, Couch GS, Morris JH, Ferrin TE (2018) UCSF ChimeraX: Meeting modern challenges in visualization and analysis. Protein Sci 27: 14–25

Gromadski KB, Rodnina MV (2004) Kinetic determinants of high-fidelity tRNA discrimination on the ribosome. Molecular cell 13: 191–200

Gutierrez E, Shin B-SS, Woolstenhulme CJ, Kim J-RR, Saini P, Buskirk AR, Dever TE (2013) eIF5A promotes translation of polyproline motifs. Molecular cell 51: 35–45

Ikeuchi K, Inada T (2016) Ribosome-associated Asc1/RACK1 is required for endonucleolytic cleavage induced by stalled ribosome at the 3’ end of nonstop mRNA. Scientific reports 6: 28234

Ikeuchi K, Tesina P, Matsuo Y, Sugiyama T, Cheng J, Saeki Y, Tanaka K, Becker T, Beckmann R, Inada T (2019a) Collided ribosomes form a unique structural interface to induce Hel2-driven quality control pathways. EMBO J 38

Ikeuchi K, Tesina P, Matsuo Y, Sugiyama T, Cheng J, Saeki Y, Tanaka K, Becker T, Beckmann R, Inada T (2019b) Collided ribosomes form a unique structural interface to induce Hel2-driven quality control pathways. The EMBO journal

Joazeiro CAP (2019) Mechanisms and functions of ribosome-associated protein quality control. Nat Rev Mol Cell Biol 20: 368–383

Juszkiewicz S, Chandrasekaran V, Lin Z, Kraatz S, Ramakrishnan V, Hegde RS (2018a) ZNF598 Is a Quality Control Sensor of Collided Ribosomes. Mol Cell 72: 469–481 e7

Juszkiewicz S, Chandrasekaran V, Lin Z, Kraatz S, Ramakrishnan V, Hegde RS (2018b) ZNF598 Is a Quality Control Sensor of Collided Ribosomes. Molecular cell 72: 469–515032704

Keedy HE, Thomas EN, Zaher HS (2018) Decoding on the ribosome depends on the structure of the mRNA phosphodiester backbone. Proc Natl Acad Sci U S A 115: E6731–E6740

Knorr AG, Schmidt C, Tesina P, Berninghausen O, Becker T, Beatrix B, Beckmann R (2019) Ribosome-NatA architecture reveals that rRNA expansion segments coordinate N-terminal acetylation. Nat Struct Mol Biol 26: 35–39

Koutmou KS, Schuller AP, Brunelle JL, Radhakrishnan A, Djuranovic S, Green R (2015) Ribosomes slide on lysine-encoding homopolymeric A stretches. Elife 4

Letzring DP, Dean KM, Grayhack EJ (2010) Control of translation efficiency in yeast by codon-anticodon interactions. RNA (New York, NY) 16: 2516–2528

Letzring DP, Wolf AS, Brule CE, Grayhack EJ (2013) Translation of CGA codon repeats in yeast involves quality control components and ribosomal protein L1. RNA (New York, NY) 19: 1208–1217

Limoncelli KA, Merrikh CN, Moore MJ (2017) ASC1 and RPS3: new actors in 18S nonfunctional rRNA decay. RNA (New York, NY) 23: 1946–1960

Ling C, Ermolenko DN (2016) Structural insights into ribosome translocation. Wiley interdisciplinary reviews RNA

Liu W, Chen C, Kavaliauskas D, Knudsen CR, Goldman YE, Cooperman BS (2015) EF-Tu dynamics during pre-translocation complex formation: EF-Tu·GDP exits the ribosome via two different pathways. Nucleic acids research 43: 9519–9528

Lu J, Deutsch C (2008) Electrostatics in the ribosomal tunnel modulate chain elongation rates. J Mol Biol 384: 73–86

Matsuo Y, Ikeuchi K, Saeki Y, Iwasaki S, Schmidt C, Udagawa T, Sato F, Tsuchiya H, Becker T, Tanaka K, Ingolia NT, Beckmann R, Inada T (2017) Ubiquitination of stalled ribosome triggers ribosome-associated quality control. Nature communications 8: 159

Murphy FVt, Ramakrishnan V (2004) Structure of a purine-purine wobble base pair in the decoding center of the ribosome. Nat Struct Mol Biol 11: 1251–2

Nakane T, Kimanius D, Lindahl E, Scheres SH (2018) Characterisation of molecular motions in cryo-EM single-particle data by multi-body refinement in RELION. Elife 7

Ozsolak F, Kapranov P, Foissac S, Kim SW, Fishilevich E, Monaghan AP, John B, Milos PM (2010) Comprehensive polyadenylation site maps in yeast and human reveal pervasive alternative polyadenylation. Cell 143: 1018–29

Pechmann S, Frydman J (2013) Evolutionary conservation of codon optimality reveals hidden signatures of cotranslational folding. Nature structural & molecular biology 20: 237–243

Pettersen EF, Goddard TD, Huang CC, Couch GS, Greenblatt DM, Meng EC, Ferrin TE (2004) UCSF Chimera--a visualization system for exploratory research and analysis. Journal of computational chemistry 25: 1605–12

Presnyak V, Alhusaini N, Chen Y-HH, Martin S, Morris N, Kline N, Olson S, Weinberg D, Baker KE, Graveley BR, Coller J (2015a) Codon optimality is a major determinant of mRNA stability. Cell 160: 1111–1124

Presnyak V, Alhusaini N, Chen YH, Martin S, Morris N, Kline N, Olson S, Weinberg D, Baker KE, Graveley BR, Coller J (2015b) Codon optimality is a major determinant of mRNA stability. Cell 160: 1111–24

Quax TE, Claassens NJ, Söll D, van der Oost J (2015) Codon Bias as a Means to Fine-Tune Gene Expression. Molecular cell 59: 149–161

Rozov A, Khusainov I, El Omari K, Duman R, Mykhaylyk V, Yusupov M, Westhof E, Wagner A, Yusupova G (2019) Importance of potassium ions for ribosome structure and function revealed by long-wavelength X-ray diffraction. Nat Commun 10: 2519

Schmidt C, Kowalinski E, Shanmuganathan V, Defenouillere Q, Braunger K, Heuer A, Pech M, Namane A, Berninghausen O, Fromont-Racine M, Jacquier A, Conti E, Becker T, Beckmann R (2016) The cryo-EM structure of a ribosome-Ski2-Ski3-Ski8 helicase complex. Science 354: 1431–1433

Schuller AP, Wu CC, Dever TE, Buskirk AR, Green R (2017) eIF5A Functions Globally in Translation Elongation and Termination. Molecular cell 66: 194–20500000

Selmer M, Dunham CM, Murphy FVt, Weixlbaumer A, Petry S, Kelley AC, Weir JR, Ramakrishnan V (2006) Structure of the 70S ribosome complexed with mRNA and tRNA. Science 313: 1935–42

Sharp PM, Li WH (1987) The codon Adaptation Index--a measure of directional synonymous codon usage bias, and its potential applications. Nucleic acids research 15: 1281–1295

Shoemaker CJ, Eyler DE, Green R (2010) Dom34:Hbs1 Promotes Subunit Dissociation and Peptidyl-tRNA Drop-Off to Initiate No-Go Decay. Science 330: 369–372

Simms CL, Kim KQ, Yan LL, Qiu J, Zaher HS (2018) Interactions between the mRNA and Rps3/uS3 at the entry tunnel of the ribosomal small subunit are important for no-go decay. PLoS genetics 14

Simms CL, Yan LL, Qiu JK, Zaher HS (2019) Ribosome Collisions Result in +1 Frameshifting in the Absence of No-Go Decay. Cell Rep 28: 1679–1689 e4

Simms CL, Yan LL, Zaher HS (2017) Ribosome Collision Is Critical for Quality Control during No-Go Decay. Molecular cell 68: 361–37300000

Tang TTL, Stowell JAW, Hill CH, Passmore LA (2019) The intrinsic structure of poly(A) RNA determines the specificity of Pan2 and Caf1 deadenylases. Nat Struct Mol Biol 26: 433–442

Tesina P, Heckel E, Cheng J, Fromont-Racine M, Buschauer R, Kater L, Beatrix B, Berninghausen O, Jacquier A, Becker T, Beckmann R (2019) Structure of the 80S ribosome-Xrn1 nuclease complex. Nat Struct Mol Biol 26: 275–280

Thanaraj TA, Argos P (1996) Ribosome-mediated translational pause and protein domain organization. Protein science: a publication of the Protein Society 5: 1594–1612

Tuller T, Waldman YY, Kupiec M, Ruppin E (2010) Translation efficiency is determined by both codon bias and folding energy. Proceedings of the National Academy of Sciences of the United States of America 107: 3645–3650

Wang J, Zhou J, Yang Q, Grayhack EJ (2018) Multi-protein bridging factor 1(Mbf1), Rps3 and Asc1 prevent stalled ribosomes from frameshifting. eLife 7

Waters MG, Blobel G (1986) Secretory protein translocation in a yeast cell-free system can occur posttranslationally and requires ATP hydrolysis. J Cell Biol 102: 1543–50

Wolf AS, Grayhack EJ (2015) Asc1, homolog of human RACK1, prevents frameshifting in yeast by ribosomes stalled at CGA codon repeats. RNA (New York, NY) 21: 935–945

Wu CC, Zinshteyn B, Wehner KA, Green R (2019) High-Resolution Ribosome Profiling Defines Discrete Ribosome Elongation States and Translational Regulation during Cellular Stress. Molecular cell

Yarus M, Folley LS (1985) Sense codons are found in specific contexts. Journal of molecular biology 182: 529–540

Yokogawa T, Kitamura Y, Nakamura D, Ohno S, Nishikawa K (2010) Optimization of the hybridization-based method for purification of thermostable tRNAs in the presence of tetraalkylammonium salts. Nucleic acids research 38

Zaher HS, Green R (2009) Fidelity at the Molecular Level: Lessons from Protein Synthesis. Cell 136: 746–762

Zhang K (2016) Gctf: Real-time CTF determination and correction. Journal of structural biology 193: 1–12

Zheng SQ, Palovcak E, Armache JP, Verba KA, Cheng Y, Agard DA (2017) MotionCor2: anisotropic correction of beam-induced motion for improved cryo-electron microscopy. Nature methods 14: 331–332

Zivanov J, Nakane T, Forsberg BO, Kimanius D, Hagen WJ, Lindahl E, Scheres SH (2018) New tools for automated high-resolution cryo-EM structure determination in RELION-3. Elife 7

