## Supplementary material for "Molecular mechanism of translational stalling by inhibitory codon combinations and poly(A) tracts": Expanded view figures

**Expanded view figures for**

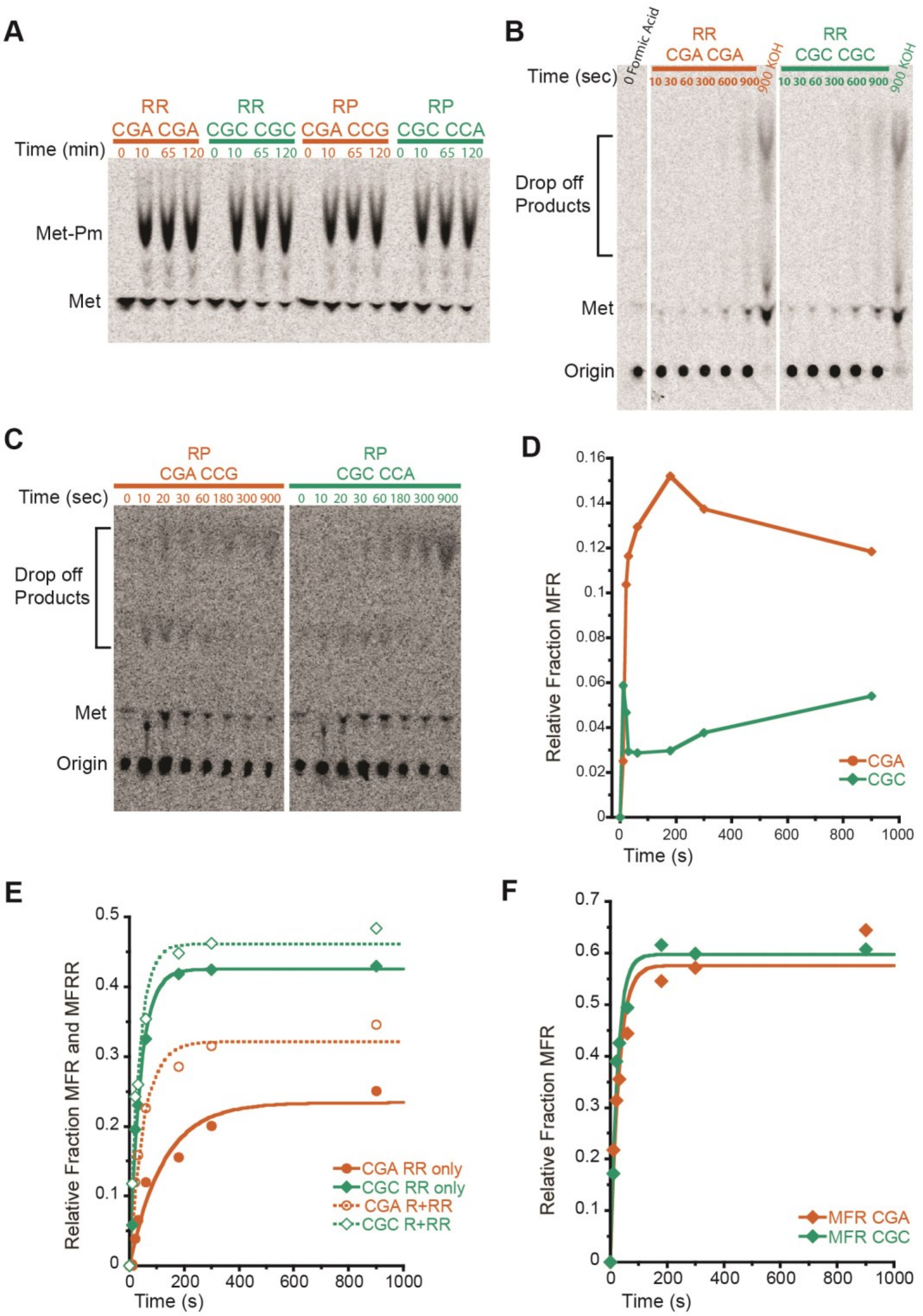


**Fig EV1: Initiation complex test of Met-Pm activity and individual product analysis of MFRR elongation.**

**(A)** Met-Pm activity for all the ICs formed with WT ribosomes on inhibitory mRNAs (red) and optimal mRNAs (green). There is no significant difference in activity at the last time point for any of the ICs. **(B)** TLC showing peptidyl tRNA drop off using the PTH assay on MFRR ICs with the inhibitory (CGA-CGA) pair (red) and the optimal (CGC-CGC) pair (green). Time points were quenched with formic acid to assess drop off and time points quenched with KOH were to monitor peptide formation as a control. There is no significant accumulation of peptidyl tRNA drop off products. **(C)** TLC showing peptidyl tRNA drop off using the PTH assay on MRPK ICs with the inhibitory (CGA-CCG) pair (red) and the optimal (CGC-CCA) pair (green).

Time points were quenched with formic acid to assess drop off. There is no significant accumulation of peptidyl tRNA drop off products. (**D**)Elongation kinetics for the MFR product within the context of MFRR elongation for the inhibitory (CGA-CGA) pair (red) and the optimal (CGC-CGC) pair (green). MFR peptide builds up on the inhibitory pair as compared to the optimal pair indicative of slow formation of the next peptide bond. **(E)** Elongation kinetics for the MFR and MFRR products together versus the final MFRR product alone for the inhibitory (CGA-CGA) pair (red) and the optimal (CGC-CGC) pair (green). The increased rate and amount of product formed for the MFR and MFRR data compared to the MFRR alone suggest that the addition of the second arginine is slower than the first. (**F**) Elongation kinetics for the addition of a single arginine MFR CGA (red) and CGC (green). The addition of the first arginine is only slightly slower for CGA again suggesting that the addition of the second arginine is the slower step.

**A**

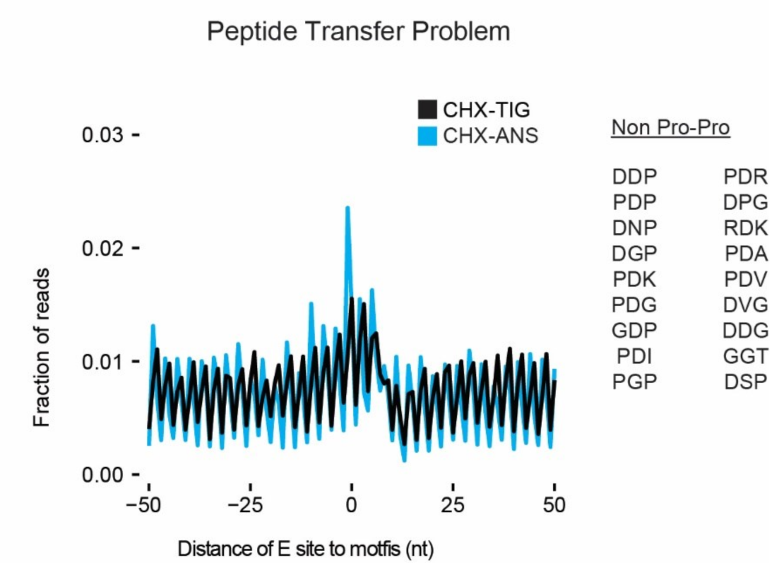


**Fig EV2: Ribosome profiling analysis showing defects in peptide bond formation.**

**(A)** Metacodon analysis of 21 nt RPFs in libraries prepared with CHX/ANS (blue) showing an increase in ribosome density at tripeptide motifs that undergo slow peptide bond formation (Schuller et al., 2017) compared to libraries prepared with CHX/TIG.

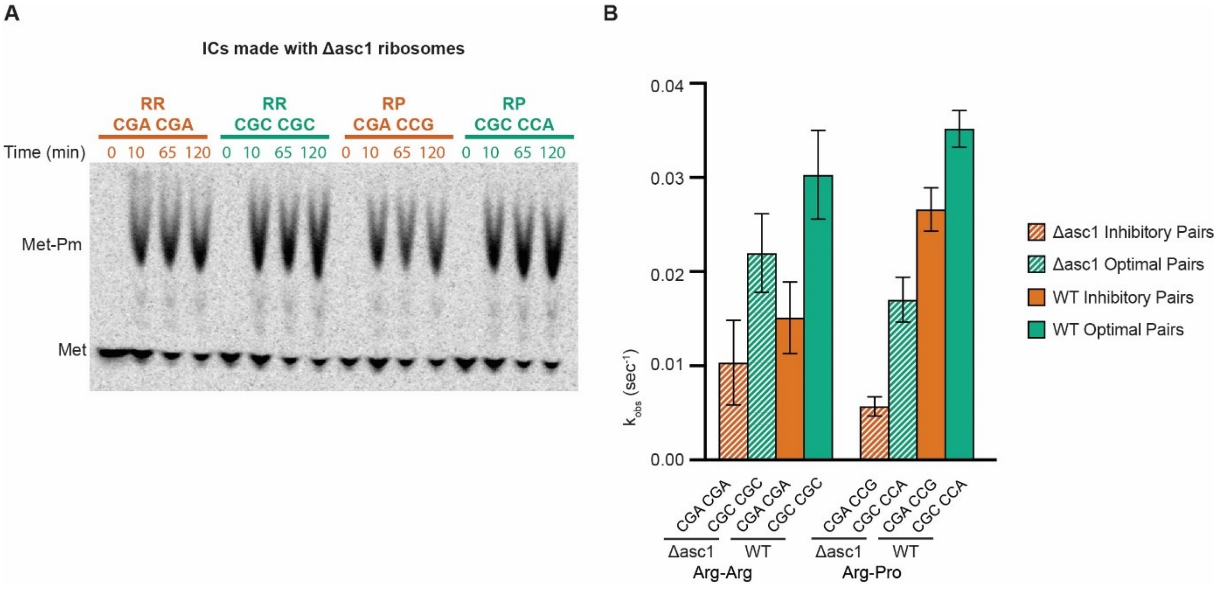


**Fig EV3: Initiation complex test of Met-Pm activity with ribosomes lacking Asc1 and their corresponding elongation rates.**

**(A)** Met-Pm activity for all the ICs formed with ribosomes from *asc1Δ* strain on inhibitory mRNAs (red) and optimal mRNAs (green). There is no significant difference in activity at the last time point for any of the ICs as compared to one another or to the Met-Pm activity for WT ICs (Figure S1A). **(B)** Comparison of observed rates of elongation at saturating tRNA concentrations for all inhibitory pairs (red) and their optimal controls (green) by ICs formed with ribosomes lacking Asc1 (hatched bars) versus WT ribosomes (solid bars). Error bars represent the standard deviation calculated from three replicate experiments with the exception of the MFRP inhibitory and optimal pairs (two replicates).

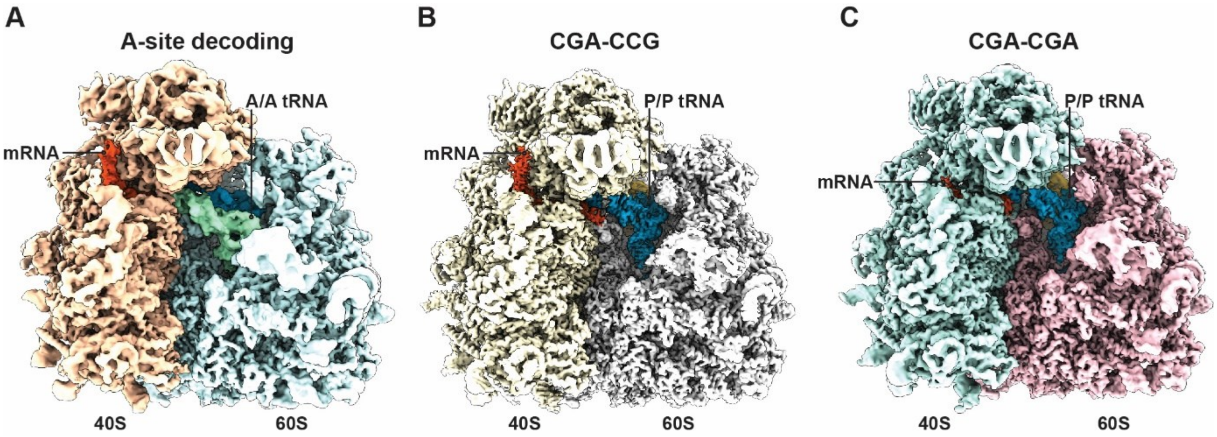


**Fig EV4: Cryo-EM structures of RNCs stalled on inhibitory codon pairs in comparison with the A site decoding situation.**

(**A-C**) Cryo-EM density maps filtered according to local resolution used to build molecular models. (**A**) Cryo-EM map of the pre-state RNC with tRNA in the A site. (**B**) Cryo-EM map of the CGA-CCG stalled RNC. (**C**) Cryo-EM map of the CGA-CGA stalled RNC.

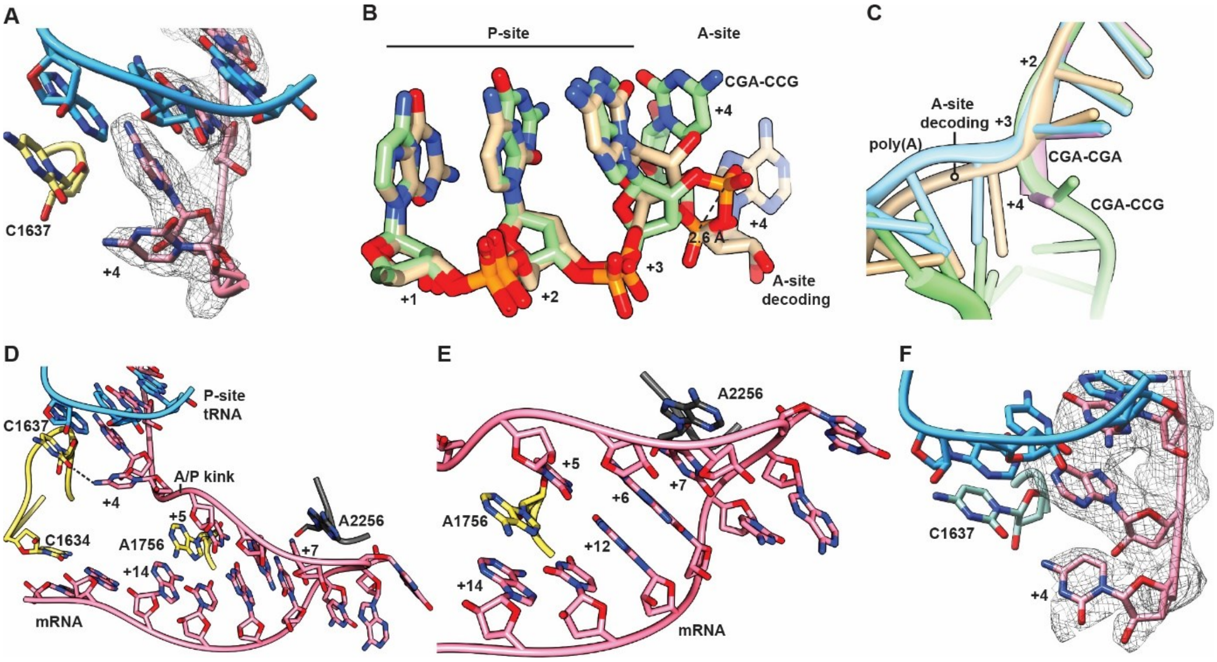


**Fig EV5: Structural details of the codon-based stalling.**

(**A**) Cryo-EM density (mesh) and stick model with cartoon phosphate backbone representing the mRNA positions +1 to +4 and their interactions in the CGA-CCG stalled ribosome. (**B**) Comparison between the A site tRNA decoding situation and the CGA-CCG stalled situation of the mRNA in positions +1 to +4. In the CGA-CCG A site, the C+4 is flipped by approximately 95° degrees towards the wobble A:I base pair and the mRNA backbone is shifted by 2.6 Å at the phosphate linking A+3 and C+4. (**C**) The effect of flipped C+4 on the general path of the mRNA in the A site. A cartoon representation of mRNAs in all four discussed 80S structures is compared. (**D**) Overview of the mRNA and it's interactions in the A site of CGA-CCG reporter stalled ribosome using stick model with cartoon phosphate backbone representation. (**E**) Detail of the tip of the hairpin from (**C**) with stabilizing stacking interactions between A2256 of the 25S rRNA and the C+7 of the mRNA and among A1756 of the 18S rRNA intercalated between the C+5 and the A+14 of the mRNA. (**F**) Cryo-EM density (mesh) and stick model with cartoon phosphate backbone representing the mRNA positions +1 to +4 and their interactions in the CGA-CGA stalled ribosome.

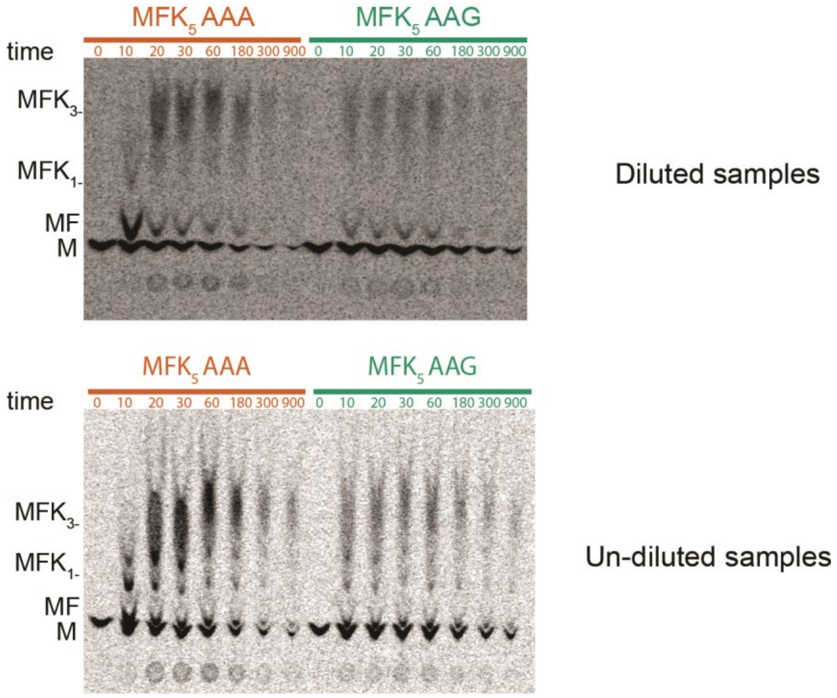


**Fig EV6: Elongation of AAA is slower than AAG on $MFK_5$ initiation complexes.**

(**A**) TLC showing peptide bond formation of $MFK_5$ messages on inhibitory AAA codons (red) and control AAG codons (green). Samples were diluted 1 to 4 µL in water (top) or undiluted (bottom). $MFK_5$ AAG complexes are making longer lysine peptides (indicated by higher bands on the TLC) than AAA at early timepoints.
