## Appendix for "Molecular mechanism of translational stalling by inhibitory codon combinations and poly(A) tracts"

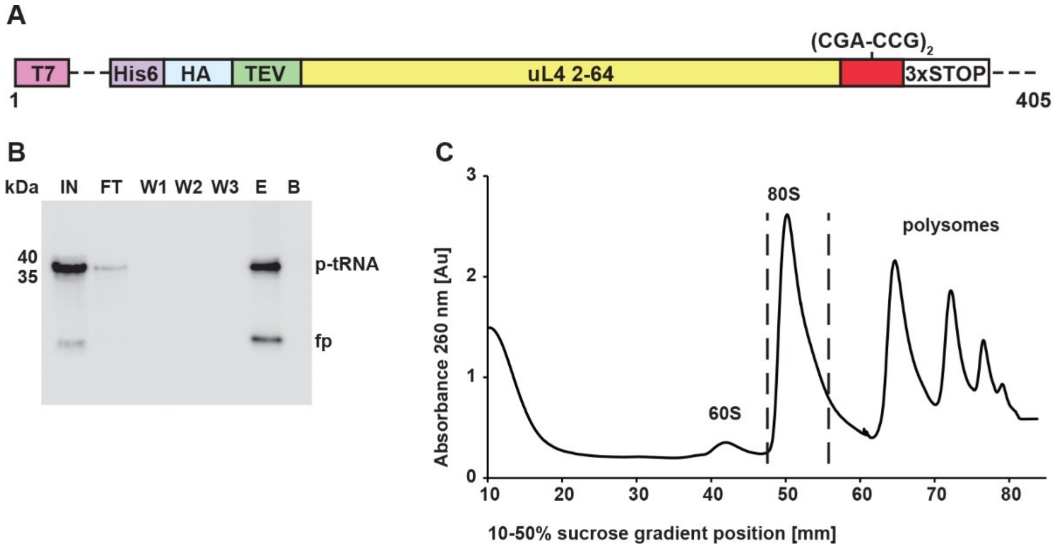


**Appendix Fig S1: CGA-CCG reporter mRNA and purification of the stalled 80S RNCs**

**A**, Schematic representation of the CGA-CCG mRNA reporter used for the structural studies. **B**, *In vitro* translation reaction (IN) using a yeast translation extract from a *ski2Δ* strain and subsequent affinity purification of His-tagged ribosome-nascent chain complexes. Fractions representing input (IN), flow through (FT), washing steps (W1 – W3), elution (E) and beads (B) were visualized by immunoblotting using anti-HA antibody. Peptidyl-tRNA (p-tRNA) and free peptide (fp) bands are indicated. **C,** The eluate was loaded on a 10-50 % sucrose gradient and fractionated. The indicated peak representing the 80S fraction was collected and concentrated using a sucrose cushion. Resuspended ribosomal pellet was used for cryo-EM sample preparation.

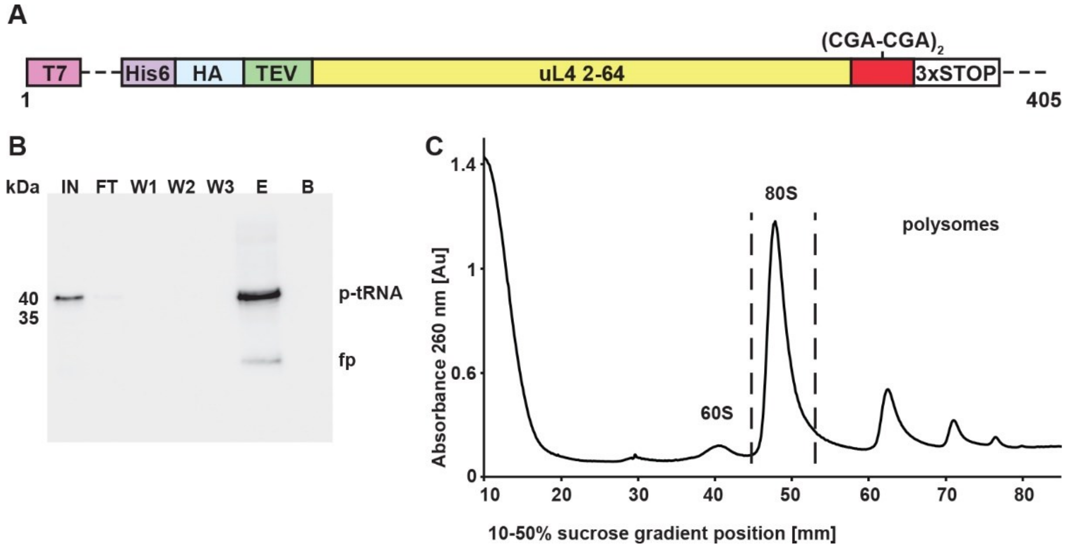


**Appendix Fig S2: CGA-CGA reporter mRNA and purification of the stalled 80S RNCs**

**A**, Schematic representation of the CGA-CGA mRNA reporter used for the structural studies. **B**, *In vitro* translation reaction (IN) using a yeast translation extract from a *ski2Δ* strain and subsequent affinity purification of His-tagged ribosome-nascent chain complexes. Fractions representing input (IN), flow through (FT), washing steps (W1 – W3), elution (E) and beads (B) were visualized by immunoblotting using anti-HA antibody. Peptidyl-tRNA (p-tRNA) and free peptide (fp) band sizes are indicated. **C,** The eluate was loaded on a 10-50 % sucrose gradient and fractionated. The indicated peak representing the 80S fraction was collected and concentrated using a sucrose cushion. Resuspended ribosomal pellet was used for cryo-EM sample preparation.

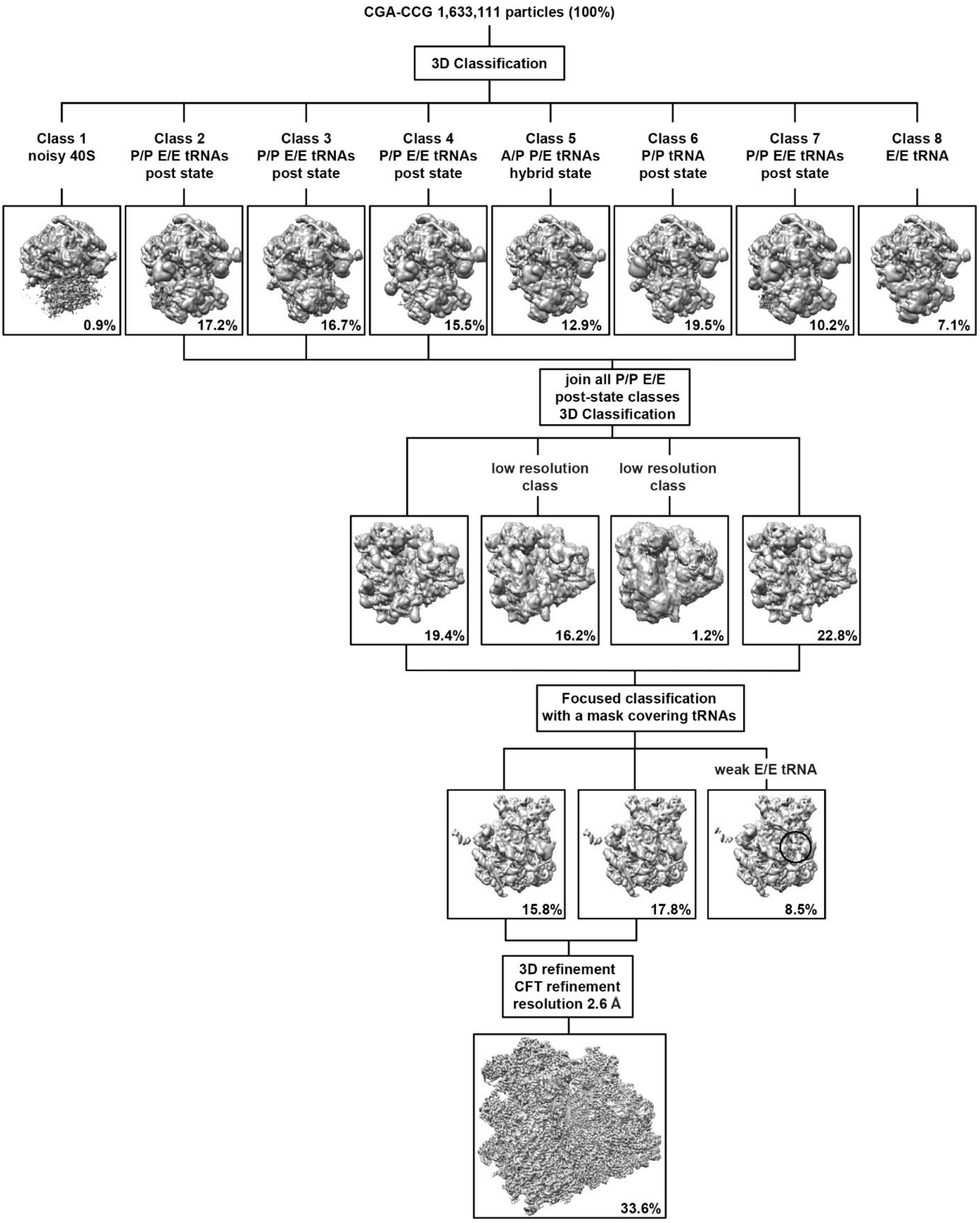


**Appendix Fig S3: 3D classification and processing scheme of the 80S ribosomes stalled on the CGA-CCG reporter mRNA**

The first 3D refined map was sorted into 8 classes. Classes 2, 3, 4 and 7 represented a vast majority of programmed ribosomal particles exhibiting the non-rotated post state with P/P and E/E site tRNAs. These classes were joined and further sub-classified, sorting out low resolution and weak E site tRNA occupancy particles. This particle category was further refined and processed as indicated (for details, see Methods).

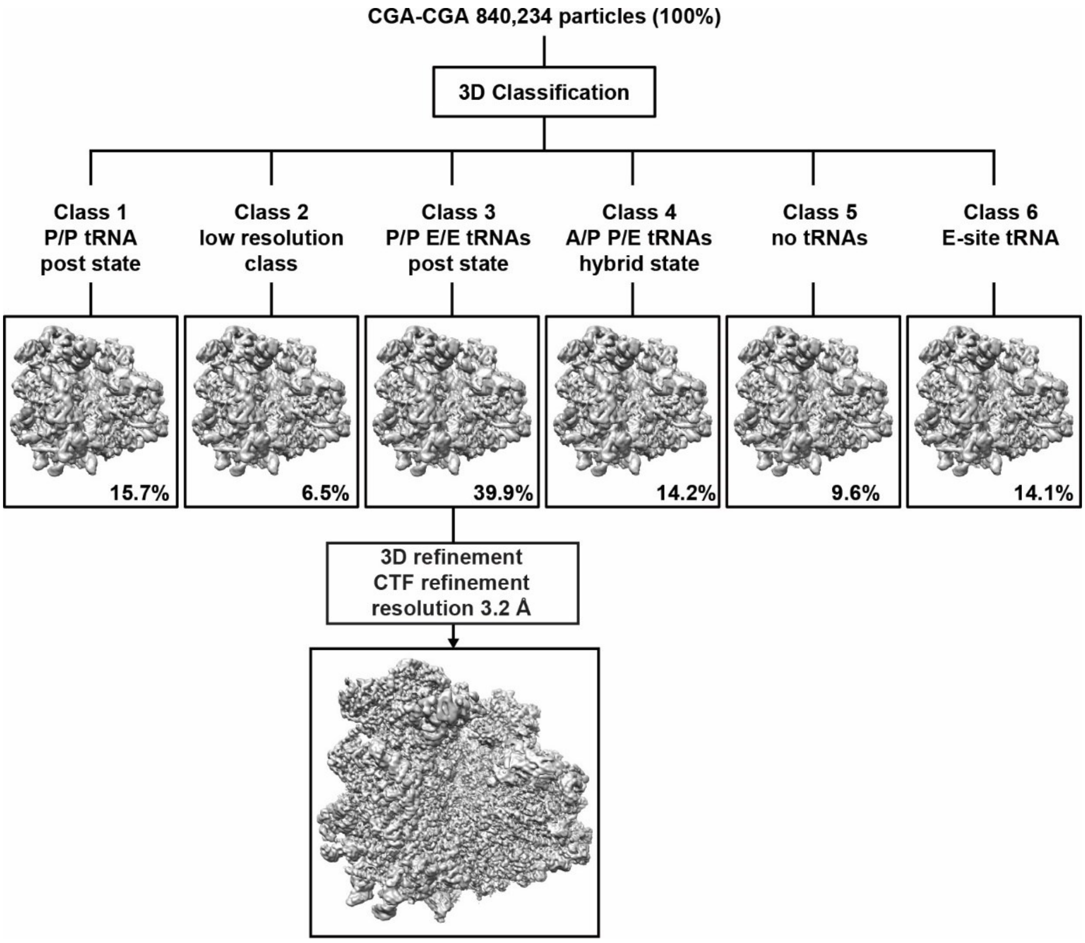


**Appendix Fig S4: 3D classification and processing scheme of the 80S ribosomes stalled on the CGA-CGA reporter mRNA**

The first 3D refined map was sorted into 6 classes. Class 3 represented a vast majority of programmed ribosomal particles exhibiting the non-rotated post state with P/P and E/E site tRNAs. This class was further refined and processed as indicated (for details, see Methods).

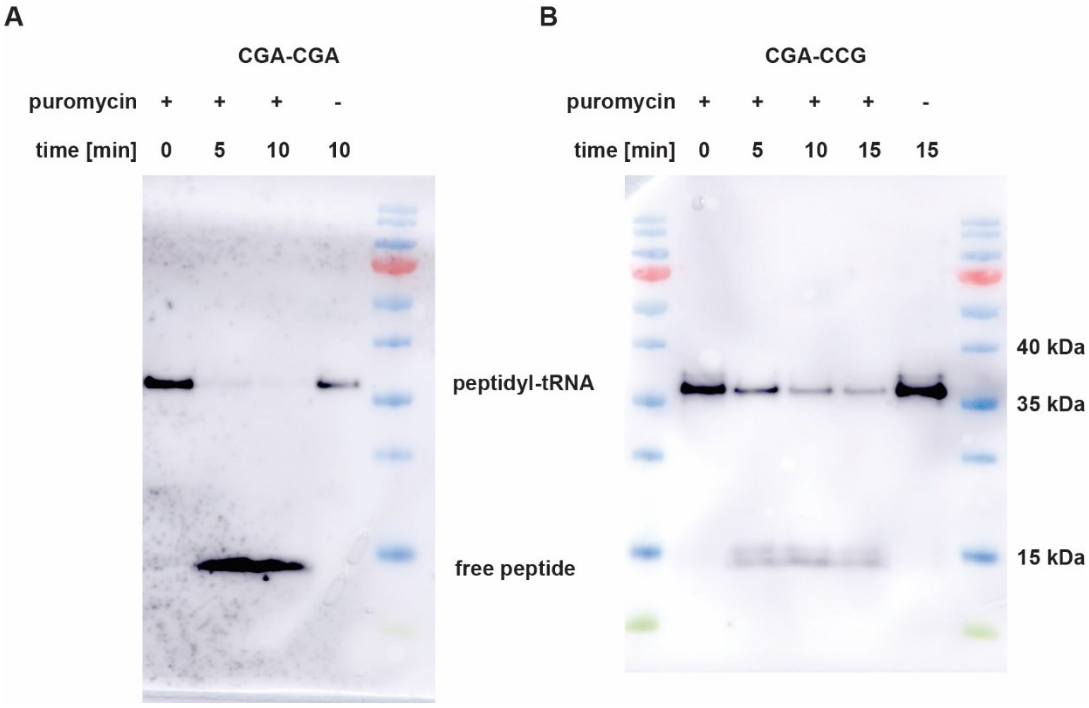


**Appendix Fig S5: Puromycin reactivity of CGA-CGA and CGA-CCG stalled RNCs**

(**A-B**) 80S fractions of RNCs isolated from sucrose density gradients (Appendix Figs 1c and 2c) were treated with 1 mM puromycin. Nascent chain species were visualized by immunoblotting using anti-HA antibody. Peptidyl-tRNA and free peptide band sizes are indicated. (**A**) 80S RNCs stalled on the CGA-CGA reporter mRNA readily reacted with puromycin releasing all detectable nascent chains within the first five minutes of the reaction. (**B**) 80S RNCs stalled on the CGA-CCG reporter mRNA reacted with puromycin slower than the CGA-CGA ones with a small fraction of unreacted peptidyl-tRNA still detectable after 15 minutes of the reaction.

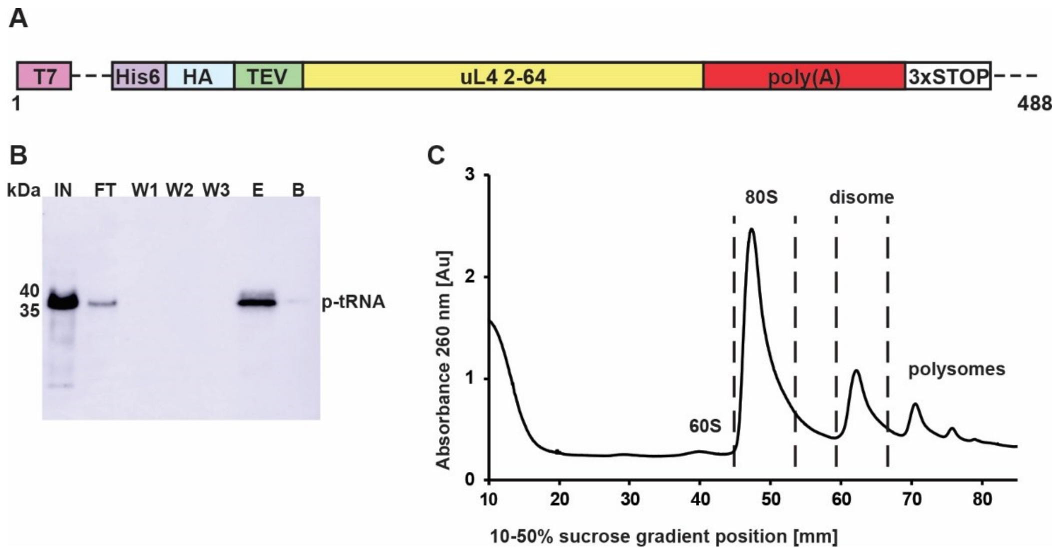


**Appendix Fig S6: Poly(A) reporter mRNA and purification of the stalled 80S RNCs**

**A**, Schematic representation of the poly(A) mRNA reporter comprising 49 consecutive adenines as a stall-inducing sequence used for the structural studies. **B**, *In vitro* translation reaction (IN) using a yeast translation extract from a *ski2Δ* strain and subsequent affinity purification of His-tagged ribosome-nascent chain complexes. Fractions representing input (IN), flow through (FT), washing steps (W1 – W3), elution (E) and beads (B) were visualized by immunoblotting using anti-HA antibody. Peptidyl-tRNA (p-tRNA) band size is indicated. **C,** The eluate was loaded on a 10-50 % sucrose gradient and fractionated. The indicated peaks representing the 80S and disome fractions were collected and concentrated using a sucrose cushion. Resuspended ribosomal pellets were used for cryo-EM sample preparation.

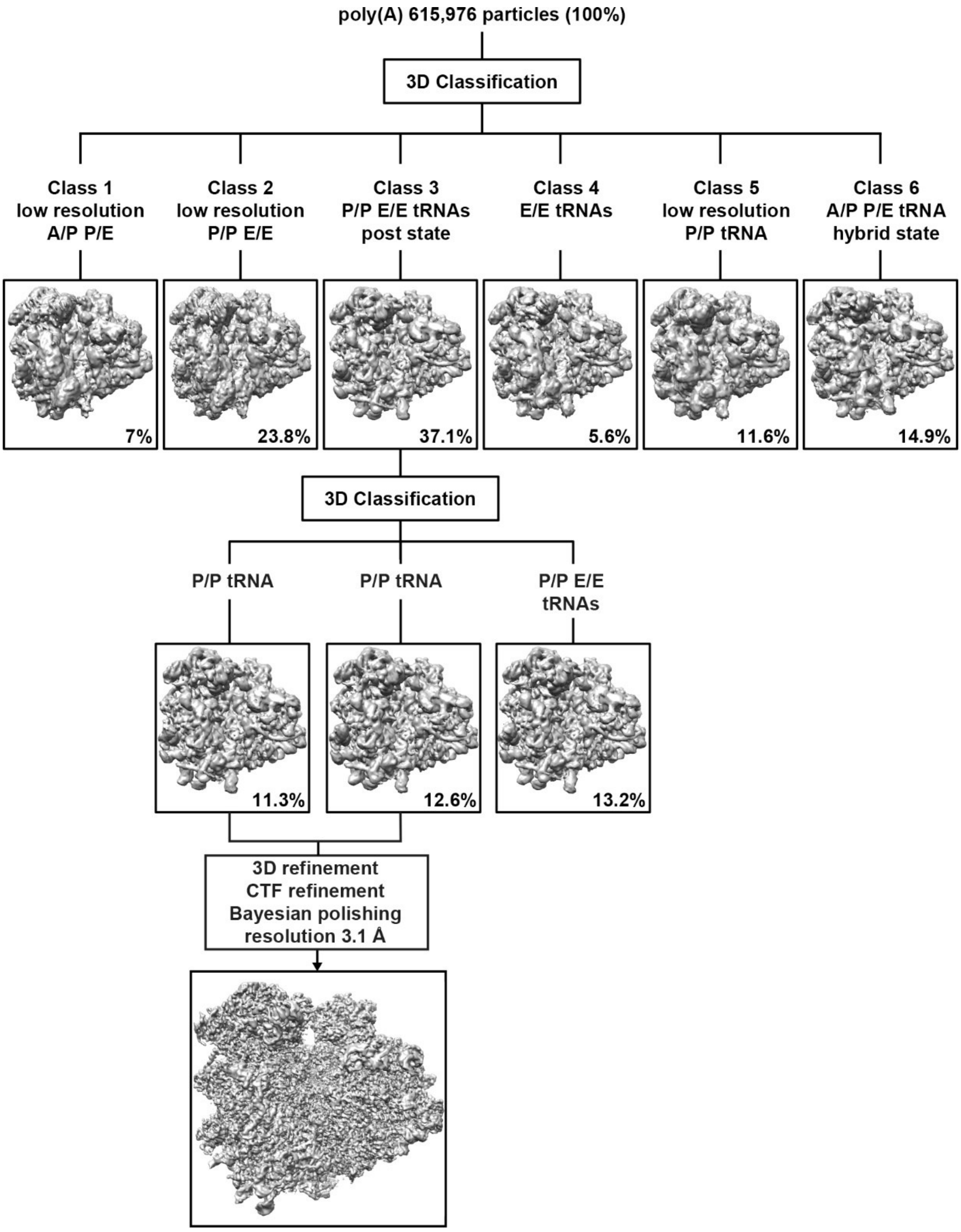


**Appendix Fig S7: 3D classification and processing scheme of the 80S ribosomes stalled on the poly(A) reporter mRNA**

The first 3D refined map was sorted into 6 classes. With the exception of classes 1 and 6 accounting for approximately 22% of particles, all other classes represented a vast majority of programmed ribosomal particles exhibiting the non-rotated post state. Class 3 was further subsorted sorting out a minor population of particles with both P/P and E/E tRNAs and two classes with P/P tRNA. These two classes were joined, resulting in a clean major population of particles with P/P tRNA. This particle category was further refined and processed as indicated (for details see Methods).

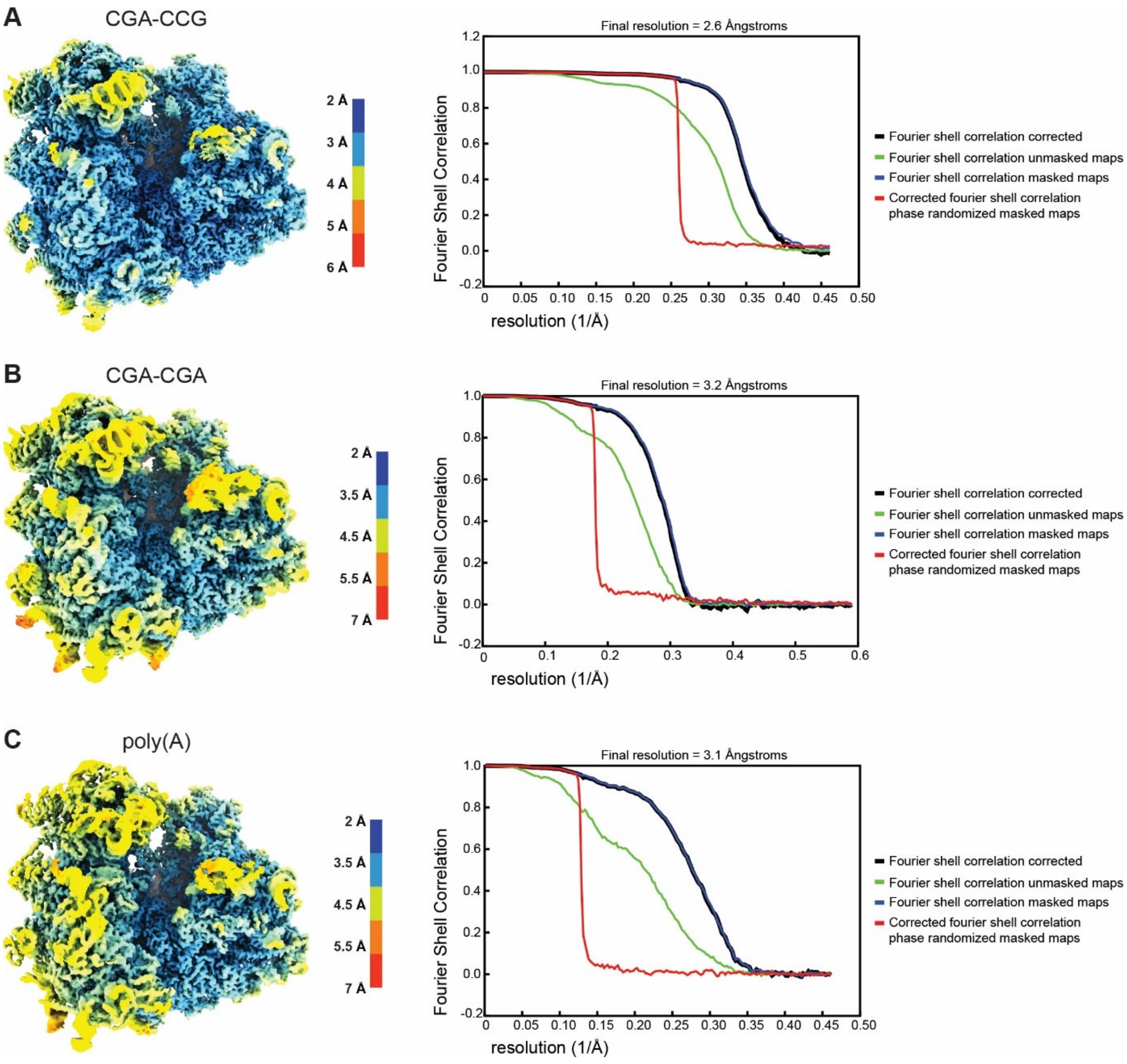


**Appendix Fig S8: Local resolution and FSC curves for the 80S cryo-EM density maps of stalled ribosomes.**

Cryo-EM density maps filtered and colored according to local resolution as estimated by Relion 3 with Fourier Shell Correlation (FSC) plots for the refined and post-processed maps of 80S ribosomes stalled on the CGA-CCG (**A**), CGA-CGA (**B**) and poly(A) (**C**) mRNA reporters.

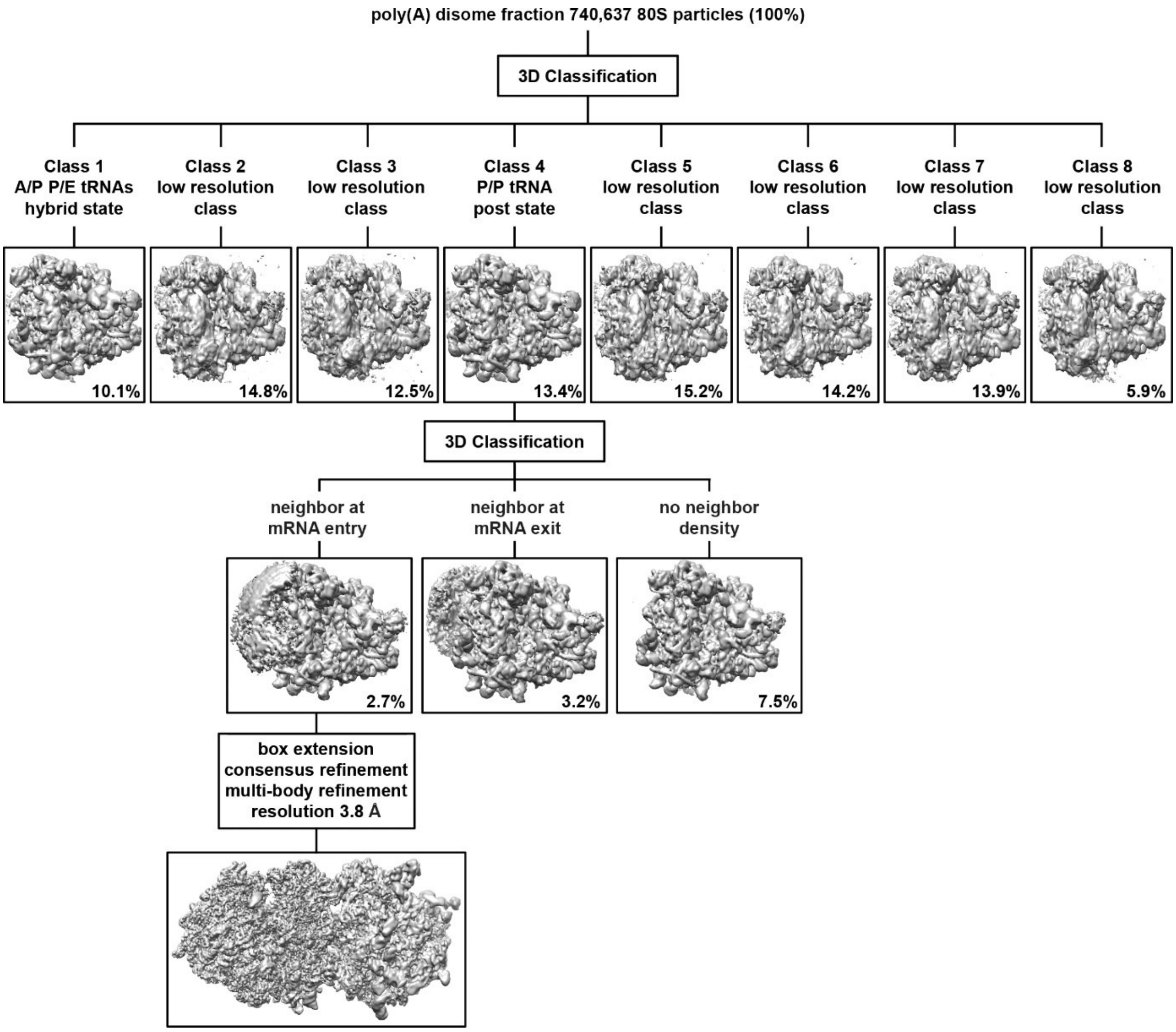


**Appendix Fig S9: 3D classification and processing scheme of the disomes stalled on the poly(A) reporter mRNA**

740,637 80S particles were 3D refined and initially separated into eight classes partly representing different translational states of the ribosome. Class 4 represented the previously characterized poly(A) stalled 80S in the non-rotated post state with P/P tRNA. This class was further sub-classified, sorting out particles with no neighboring ribosome and revealing two subclasses with approximately the same share of particles. These two classes represented the first stalling and the second colliding ribosome judging by the density of the neighbor ribosome. Further processing of the first stalling post state ribosome (with neighbor density at mRNA exit) yielded a standard post-hybrid disome assembly. However, further processing of the second colliding ribosome in the post state (with neighbor density at mRNA entry) revealed a novel post-post disome assembly. The indicated processing procedure is described in more detail in Methods.

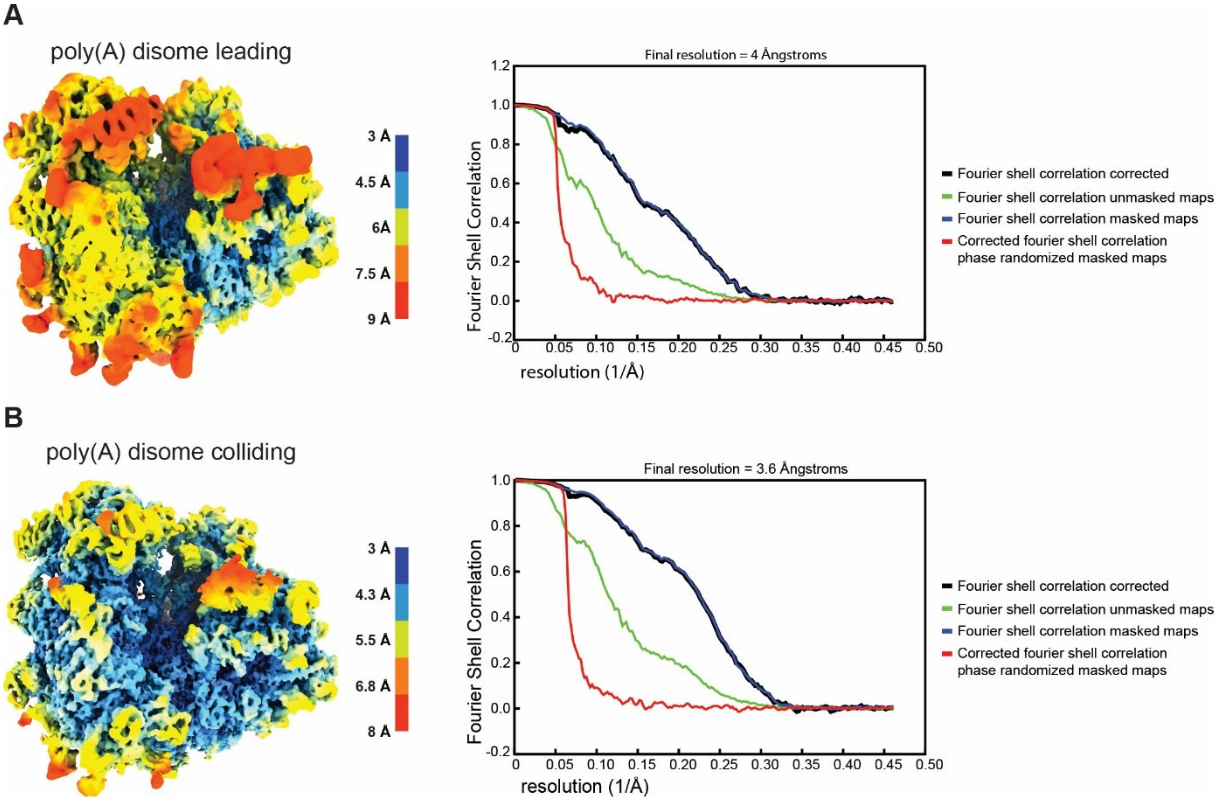


**Appendix Fig S10: Local resolution and FSC curves for the individually refined 80S cryo-EM density maps of poly(A) stalled disomes.**

Cryo-EM density maps filtered and colored according to local resolution as estimated by Relion 3 with Fourier Shell Correlation (FSC) plots for the individually refined and post-processed maps of the first stalling (**A**) and second colliding (**B**) ribosomes stalled on the poly(A) mRNA reporter.
